# A comprehensive map of the dendritic cell transcriptional network engaged upon innate sensing of HIV

**DOI:** 10.1101/579920

**Authors:** Jarrod S. Johnson, Nicholas De Veaux, Alexander W. Rives, Xavier Lahaye, Sasha Y. Lucas, Brieuc Pérot, Marine Luka, Lynn M. Amon, Aaron Watters, Alan Aderem, Nicolas Manel, Dan R. Littman, Richard Bonneau, Mickaël M. Ménager

## Abstract

Transcriptional programming of the innate immune response is pivotal for host protection. However, the transcriptional mechanisms that link pathogen sensing with innate activation remain poorly understood. During infection with HIV-1, human dendritic cells (DCs) can detect the virus through an innate sensing pathway leading to antiviral interferon and DC maturation. Here, we developed an iterative experimental and computational approach to map the innate response circuitry during HIV-1 infection. By integrating genome-wide chromatin accessibility with expression kinetics, we inferred a gene regulatory network that links 542 transcription factors with 21,862 target genes. We observed that an interferon response is required, yet insufficient to drive DC maturation, and identified PRDM1 and RARA as essential regulators of the interferon response and DC maturation, respectively. Our work provides a resource for interrogation of regulators of HIV replication and innate immunity, highlighting complexity and cooperativity in the regulatory circuit controlling the DC response to infection.

**Graphical Abstract:** 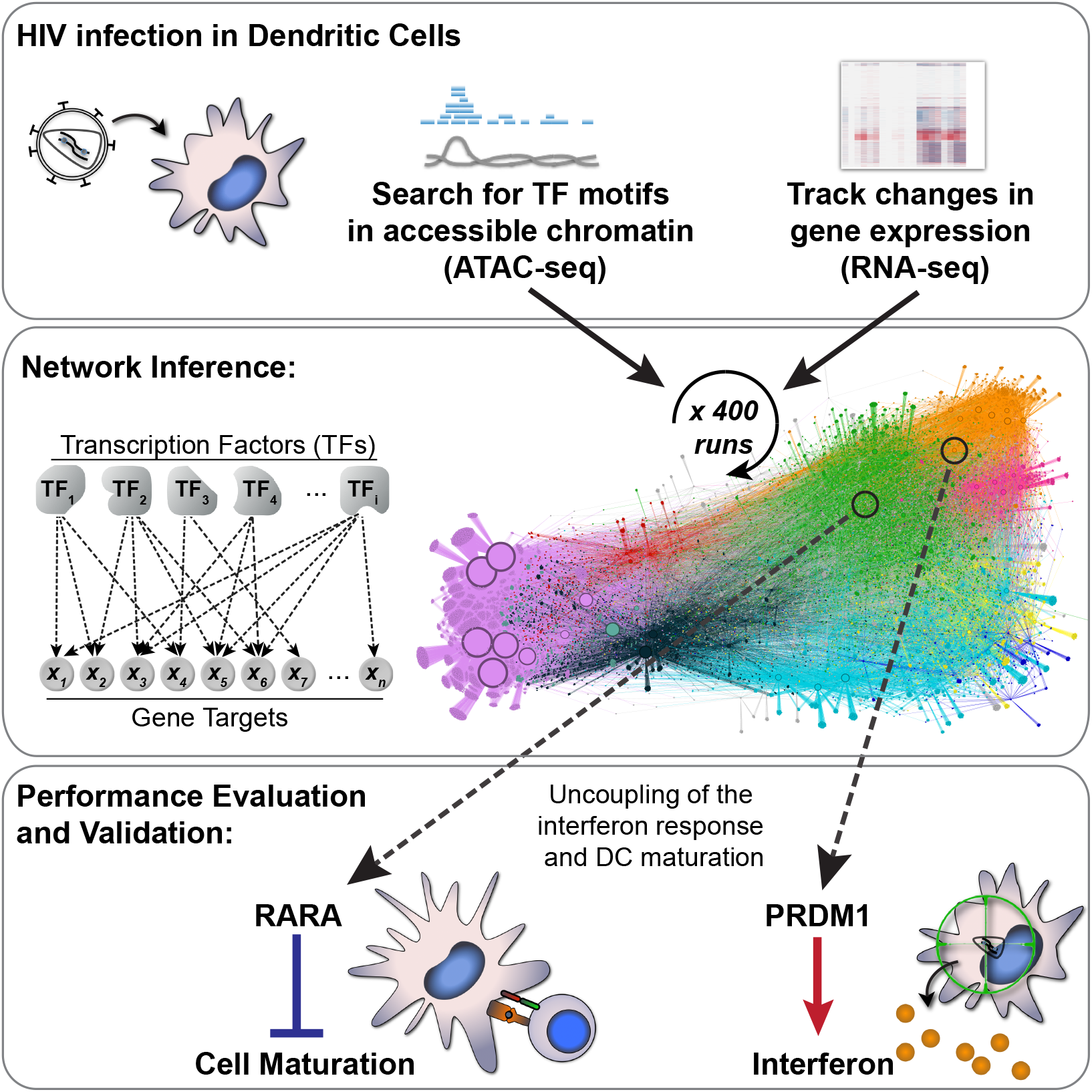

## Introduction

The host’s ability to rapidly alter gene expression in order to defend against infection is a central element of the innate immune response. Host-encoded pattern recognition receptors (PRRs) detect components of foreign microorganisms and self-derived immunostimulatory products (Cao, 2016; Ivashkiv and Donlin, 2014; Iwasaki and Medzhitov, 2015). When a pathogen is sensed, PRRs initiate signal transduction cascades that lead to activation of multiple transcription factors (TFs), which subsequently rewire gene expression to protect the host. Considering that aberrations in innate immunity are hallmarks of many disorders, including chronic viral diseases, neurodegeneration, diabetes, and cancer (Corrales et al., 2017; Heneka et al., 2014; Wada and Makino, 2016), it is not surprising that transcriptional activation of innate immune signaling is under tight control, with the goal of maintaining a sensitive response to infectious threats while avoiding unwanted inflammation and autoimmunity. In the case of HIV-1 infection, however, innate immune responses are insufficient for host protection and become dysregulated during progression to AIDS (Fernandez et al., 2011; Sandler et al., 2014; Schoggins et al., 2011).

Dendritic cells (DCs) serve key functions in host defense and are among the first cells thought to contact HIV-1 during transmission (Iijima et al., 2008). Myeloid DCs express an arsenal of PRRs and link innate detection of microbes to activation of pathogen-specific adaptive immune responses (Banchereau et al., 2000; Thery and Amigorena, 2001). These cells express cell surface receptors for HIV-1 entry, but the virus undergoes limited productive infection in DCs and does not trigger robust immune responses due to the presence of restriction factors (Granelli-Piperno et al., 2004; Manel et al., 2010; Smed-Sorensen et al., 2005). The primary restriction factor in myeloid DCs is SAMHD1, an enzyme that exhibits phosphohydrolase activity and depletes the cellular pool of dNTPs required for HIV reverse transcription (Hrecka et al., 2011; Laguette et al., 2011). This restriction can be overcome if DCs are first exposed to virus-like particles that deliver the lentiviral accessory protein, Vpx (absent in HIV-1 but encoded by SIV and HIV-2) (Goujon et al., 2006; Mangeot et al., 2000). Vpx targets SAMHD1 for degradation, enabling productive HIV-1 infection, sensing of viral components, and activation of innate immune responses (Manel et al., 2010).

Innate immune responses against HIV-1 are triggered in myeloid DCs by the sensor cyclic GMP-AMP synthase (cGAS), which detects reverse-transcribed HIV cDNA in a process that requires concomitant HIV capsid protein interaction with the cellular protein NONO, and is facilitated by other proximal factors (Gao et al., 2013; Jonsson et al., 2017; Lahaye et al., 2018; Yoh et al., 2015). Downstream of innate sensing initiated by cGAS, several transcription factors are activated, including IRF and NF-κB family members, which drive induction of interferons (IFNs), IFN-stimulated genes (ISGs), and inflammatory cytokines, and promote DCs to transition from an inactive immature state to a mature, activated state. In addition to upregulating innate antiviral factors, mature DCs express at their cell surface the costimulatory factors CD80 and CD86, which are critical for programming adaptive responses (Goubau et al., 2013; Iwasaki, 2012). The DC transcriptional response circuitry involves feedback loops that engage multiple activator and repressor TFs that collectively influence thousands of gene targets during IFN signaling and DC maturation (Ivashkiv and Donlin, 2014). For these reasons it is difficult to understand the constellation of TF-target gene connections that operate during innate immune responses using traditional approaches.

Work from our groups and others has demonstrated that gene regulatory network inference, when applied to study dynamic systems such as macrophage activation, Th17 lymphocyte polarization, and the innate immune response to cytosolic DNA, has predicted the functions of key transcriptional regulators whose involvement was previously unknown (Ciofani et al., 2012; Gilchrist et al., 2006; Lee et al., 2013; Ramsey et al., 2008). Our earlier computational methods pioneered the use of time series perturbations and the incorporation of structured “prior information” into gene regulatory network inference (Bonneau et al., 2006; Greenfield et al., 2013; Greenfield et al., 2010; Madar et al., 2010). In this report we have demonstrated that network inference is improved by ensemble-learning across hundreds of individual computational runs, with each run predicated on subsampled information in the “prior” network. We have integrated chromatin accessibility data together with genome-wide measurements of gene expression to infer and experimentally validate a network describing the human dendritic cell transcriptional circuitry that is engaged upon HIV sensing.

## Results

### Perturbation of human DCs in face of diverse innate immune and viral stimuli

With the goal of better understanding how DCs respond to innate immune stimuli, we exposed immature monocyte-derived DCs to a battery of innate immune agonists and viral challenges, generating data-sets comprised of RNA-sequencing (RNA-seq) and assays for transposase-accessible chromatin (ATAC-seq) (Figure 1A; Table S1). We tracked the kinetics of gene expression and DC maturation during HIV-1 infection compared to classic innate agonists by infecting DCs with a single-cycle HIV-1 reporter virus (HIV-1-GFP), transducing with a non-replicating lentivirus (LKO-GFP), or stimulating in parallel with the TLR4 agonist, lipopolysaccharide (LPS) or the double-stranded RNA mimetic, polyinosinic:polycytidylic acid (pIC) for 2, 8, 24, and 48 h. LPS and pIC triggered rapid changes in gene expression in human DCs across several gene clusters (Figure 1B and Table S2), resembling what has been reported for mouse bone marrow-derived DCs (Amit et al., 2009; Chevrier et al., 2011). Similarly, infection with HIV-1-GFP led to induction of innate immune genes (Figure 1B; clusters 5 & 8), but did so with delayed kinetics compared to LPS and pIC, likely due to time-dependent accumulation of reverse transcription products, integration, and virus replication progressing over the first 24 h (Gao et al., 2013; Johnson et al., 2018). Gene expression profiles were consistent with the timing of DC maturation as scored by flow cytometry (Figure S1A), with LPS and pIC stimulation leading to early and robust induction of CD86. DCs infected with HIV-1-GFP did not mature until 48 h, and only minimally responded to LKO-GFP (Figure S1A), as we previously described (Johnson et al., 2018; Manel et al., 2010).

**Figure 1.**
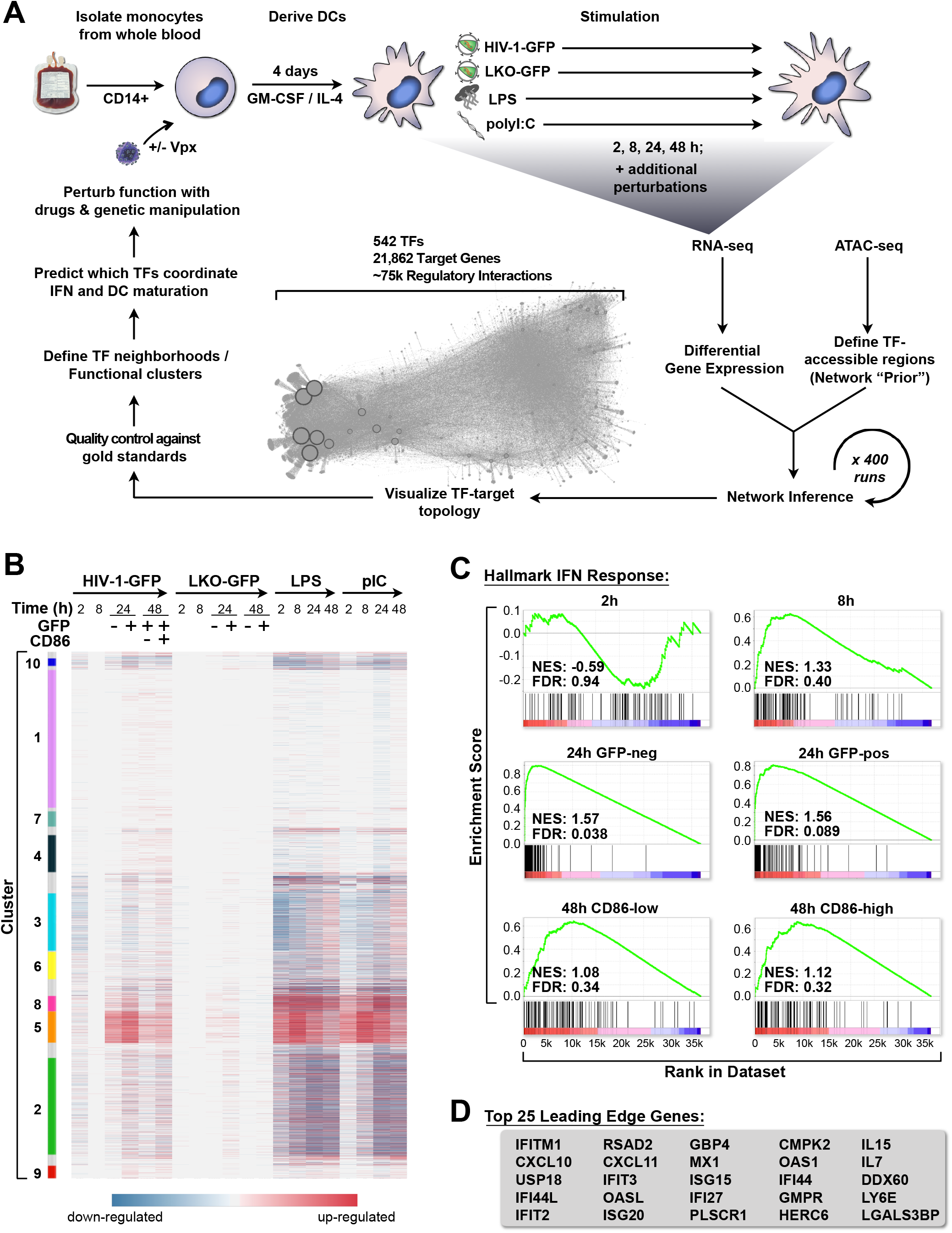
HIV-1 infection triggers a type I IFN response in DCs. (A) Schematic of the human DC network project depicting derivation of monocyte-derived DCs, time series stimulation with HIV-1-GFP, LKO-GFP, LPS, and polyI:C, followed by network inference, network visualization, *in silico* quality control, and experimental validation. (B) Heat map of differentially expressed genes in DCs across time (2, 8, 24 and 48 h), selected at FDR < 0.1 and sorted by Louvain modularity cluster (see STAR Methods). (C) GSEA plots for the Hallmark IFN Response in DCs for all described HIV-1-GFP time points compared to mock, indicating Normalized Enrichment Score (NES) and False Discovery Rate (FDR). Enrichment is considered significant at FDR < 0.25. (D) Leading edge genes from GSEA for HIV-1-GFP infections at 24 h. See also Figure S1 and Tables S1-2.

In agreement with the earlier analyses, we observed that RNA-seq samples from infected and uninfected DCs could be clearly distinguished when visualized by Principal Component Analysis (PCA) (Figure S1B). Samples at 24 and 48 h time points could be further separated when grouped specifically by time or cell sorting condition: GFP-negative (HIV-exposed but not expressing GFP), GFP-positive (HIV-infected), HIV-CD86-low (HIV-infected, immature DC), and HIV-CD86-high (HIV-infected, mature DC) (Figure S1C). We also used gene set enrichment analysis (GSEA) (Subramanian et al., 2005) to evaluate the qualitative nature of the innate response and uncovered strong associations between HIV-infected samples and gene sets for IFN alpha/beta signaling, inflammatory signaling, and DC maturation (full GSEA results available in supplemental materials). The most significant enrichment was found with the Hallmark IFN Response (Figure 1C), which peaked at 24 h post infection. Canonical ISGs were highly upregulated during HIV-1-GFP infection (Figure 1D) and their induction correlated with activation of *IFNB1* and *IFNL1* (Figure S1D). Having characterized the DC transcriptional response to HIV-1 infection, which included known maturation and interferon response signatures, we next sought, to identify regions of open chromatin that may be accessible to TF binding in order to define possible TF-to-gene target relationships that regulate the innate response.

### ATAC-seq reveals time-dependent chromatin opening at innate immune gene promoters

Analyzing genome-wide chromatin accessibility represents a powerful way to assess the presence of regulatory elements such as promoters and enhancers in mammalian cells (Buenrostro et al., 2013; Johnson et al., 2018). To match the experimental conditions used for RNA-seq, we profiled changes in open chromatin by performing ATAC-seq on DCs that were mock treated or infected with HIV-1-GFP for 2, 8, 24, and 48 h, and sorted by flow cytometry (Figure 2A-B). We identified 88k high-confidence peaks across all ATAC-seq conditions (Table S3). Similar to the RNA-seq samples, the ATAC-seq samples for mock and HIV-1-GFP could be separated by PCA through time and status of infection with minimal variation between donors (Figure S2A-B). By plotting genome-wide changes in gene expression together with changes in chromatin accessibility at transcription start sites, we observed that global changes in chromatin accessibility are temporally linked with stages of HIV-1 single-round infection and DC maturation (Figure 2C). At 24 h post infection, we found that the induction of *IFNB1, IFNL1*, and most Hallmark IFN response genes in GFP-negative and GFP-positive populations were linked with increases in chromatin accessibility. By 48 h, expression intensity and chromatin accessibility of these genes began to subside, with particularly noticeable reductions in chromatin accessibility for *IFNB1* and *IFNL1* to levels below baseline.

**Figure 2.**
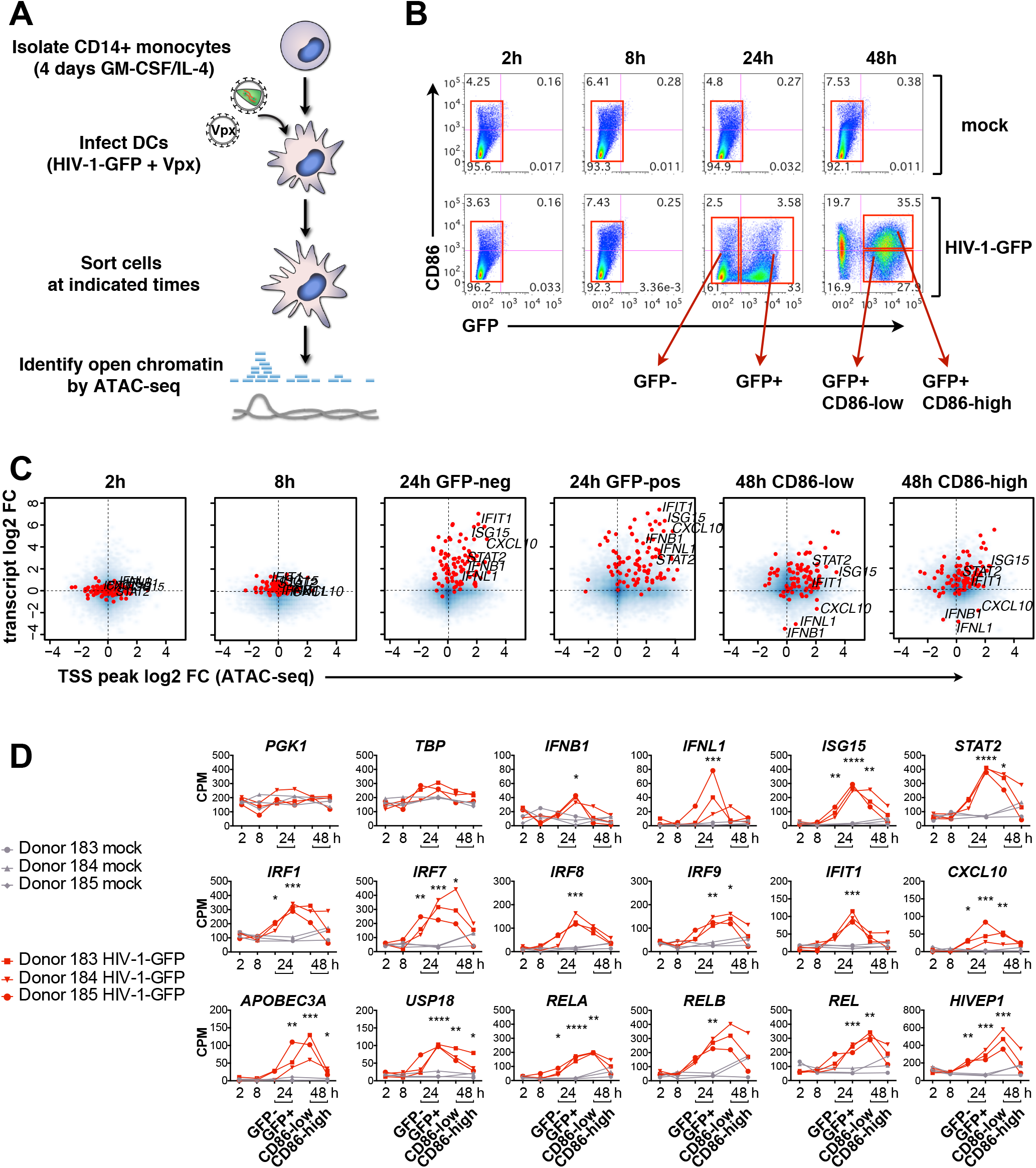
Transient changes in chromatin accessibility correspond to activation of the innate response during HIV infection. (A) Illustration depicting ATAC-seq sample processing (B) Flow cytometry plots of DCs sorted (red boxes) for ATAC-seq after infection with HIV-1-GFP at 2, 8, 24, and 48 h (MOI = 5). Plots show CD86 vs GFP expression and are representative data from 1 of 3 donors. (C) Smooth scatter density plots of HIV-1-GFP infected DCs from sorted populations as in (B) showing the genome-wide relationship between transcript levels (RNA-seq average log2 fold change) TSS chromatin accessibility (ATAC-seq average log2 fold change). Blue: density of points across the entire genome. Red: Hallmark IFN response genes as in Figure 1C. The position of *IFNB1, IFNL1*, and select ISGs are labeled for each condition. (D) ATAC-seq gene-associated peak values shown as counts per million reads (CPM) for the indicated time points and conditions. Lines connect samples from independent donors. Statistics were calculated on log2-transformed data using a 2-way ANOVA with multiple comparisons; * p < 0.05; ** p < 0.01; *** p < 0.001; **** p < 0.0001. See also Figure S2 and Table S3.

To investigate changes in chromatin accessibility in more detail, we plotted promoter-associated ATAC-seq peak height across the time series for two housekeeping control genes (*PGK1* & *TBP*), the interferon genes *IFNB1 and IFNL1*, well-defined ISGs, and IRF and NF-κB family members. Maximum chromatin accessibility for *IFNB1, IFNL1* and several ISGs was detected at 24 h post infection in cells expressing the GFP reporter (Figures 2D; S2C-D). For IFN-related genes and the NF-κB family members, *RELA, RELB, REL*, and the related TF, *HIVEP1*, chromatin accessibility was higher in HIV-infected, CD86-low DCs that were not fully mature as compared to CD86-high, mature DCs (Figure 2D), suggesting that the chromatin state is linked to DC maturation status.

To determine whether mapping open chromatin using ATAC-seq in HIV-infected DCs offered specific advantages over publicly available datasets, we compared our ATAC-seq peaks to steady-state open chromatin data from CD14+ monocytes (Figure S2C-D). Several ISGs displayed high chromatin accessibility at baseline (*STAT1, STAT2, IRF1, IRF7, IRF9, OASL, HLA-C*, and *ISG20*), with peaks detected in both DC ATAC-seq samples and CD14+ monocyte data available from ENCODE (Figure S2C). HIV-1-GFP infection led to a transient increase in chromatin accessibility at ISG promoters that corresponded with known binding sites for IRF3, STAT1, and NF-κB (RELA). In contrast, *IFNB1*, *IFNL1*, and other ISGs (*CXCL10, CXCL11, ISG15, LY6E, USP18*, and *IFIT1*), displayed open chromatin in DCs infected with HIV-1-GFP, but not in mock-treated cells or in CD14+ monocytes (Figure S2D), supporting the use of cell- and condition-specific ATAC-seq data as a basis for network inference.

### Inferred network of DC transcriptomic changes following innate immune responses

To infer a predictive gene regulatory network we adapted our previously published algorithm (the Inferelator), which was designed to learn from mixes of steady state and dynamic data (Bonneau et al., 2006). The method can incorporate multiple data types to influence network model selection, including TF occupancy, cooperativity, and transcriptional profiling in response to innate immune stimuli (Figures 1 & 2). High-throughput methods like ATAC-seq and ChIP-seq can be used to guide network inference by defining “prior” information on network architecture, dramatically improving network model selection and predictive power (Arrieta-Ortiz et al., 2015; Miraldi et al., 2019; Siahpirani and Roy, 2017). Key to this study, we extended our previously published computational methods for learning the regulation of gene networks through the Inferelator (Arrieta-Ortiz et al., 2015; Bonneau et al., 2006; Ciofani et al., 2012; Greenfield et al., 2010; Madar et al., 2010) by integrating results obtained from RNA-seq and ATAC-Seq experiments performed at the bulk level in a time-course fashion (Figure 3A).

**Figure 3.**
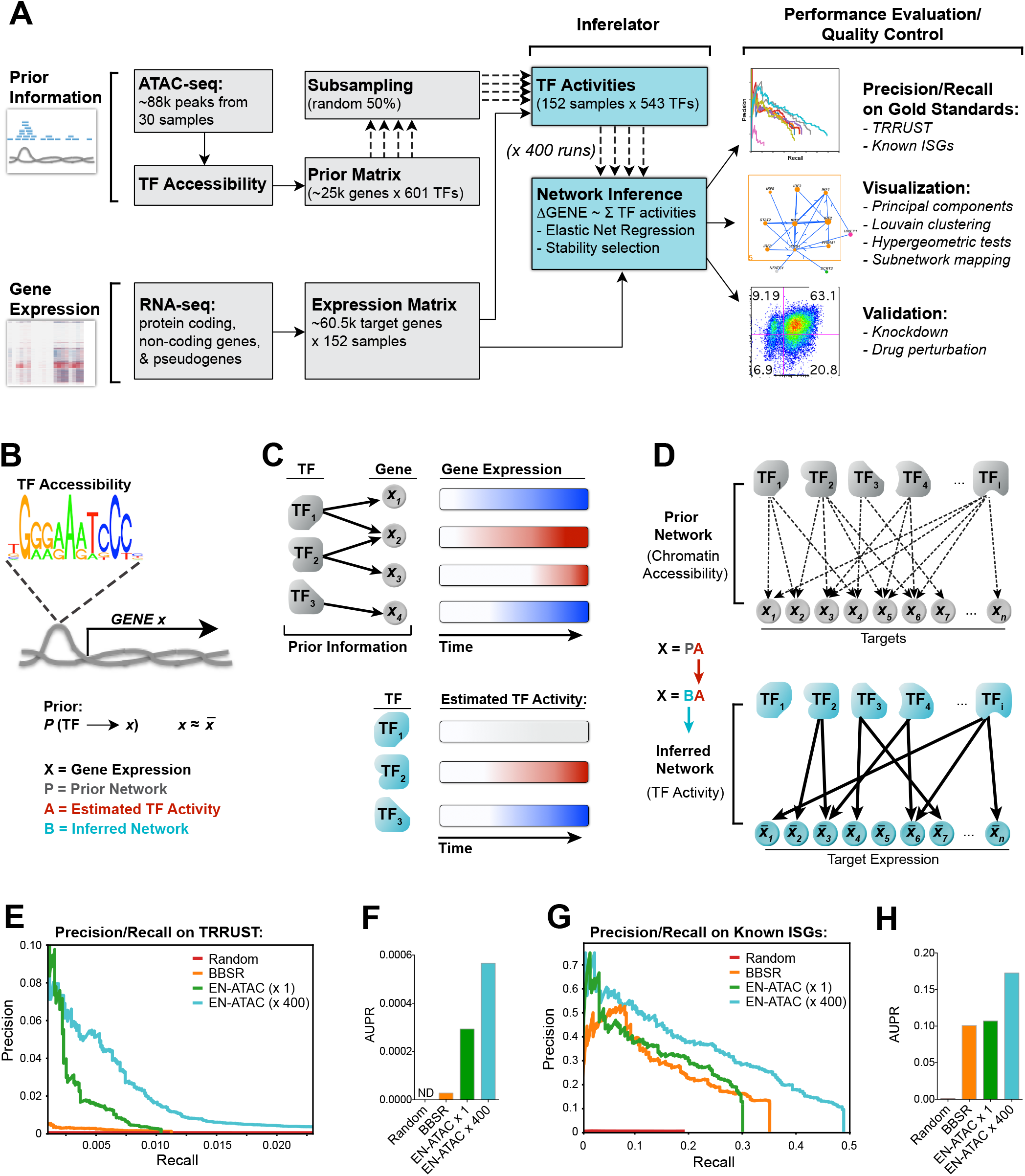
Repeated subsampling of prior information improves network inference. (A) Pipeline for network inference. Prior information (TF accessibility) and gene expression time series data were used as inputs for the Inferelator. Output networks underwent performance evaluation and quality control as indicated. (B) The prior matrix denotes a possible connection between a TF and gene X if that TF’s binding motif is found in accessible chromatin upstream (−1kb) or within gene X. (C) Schematic illustrating how TF activity is estimated by summing predicted target expression over time (red = increase; blue = decrease). (D) TF and gene target connections in the prior network are pruned using an Elastic Net regression to yield the final inferred network. (E) Precision/Recall plots and area under precision/recall (AUPR) curves (F) for the indicated networks scored against the TRRUST database (see STAR Methods). (G) Precision/Recall plots and AUPR curves (H) for the indicated networks scored against known ISGs (Schoggins et al., 2011). See also Figure S3 and Table S4.

Here, prior information on network architecture is derived from the 88k peaks of chromatin accessibility data generated by our ATAC-seq experiments that tracked DC responses to HIV infection (Figures 2A & 3A). Using curated TF binding motif databases, we established possible TF-gene target relationships by searching ATAC-seq peaks for TF motifs that we located within or up to 1 kb upstream of gene bodies (Figure 3B). This information was used to build a prior matrix that connected each gene with its possible TF regulators (Table S4). We then estimated TF activity over the course of stimulation based on the combined expression of a TF’s predicted gene targets (Figure 3C). This step approximates the effect of unmeasured parameters, such as post-translational regulation and protein-protein interactions, on TF activity in a condition-dependent manner. We were thus able to model changes in TF activity that are semi-independent of changes in TF expression, as observed for IRF3 (Figure S3A, S3B). IRF3 activity was predicted to increase in response to LPS, pIC and HIV sensing, which are all known to drive IFN production through IRF3 phosphorylation. Our inference model also predicted that activity and expression for STAT2, IRF7, and RELA correlated with innate stimulation, as expected, given their well-defined roles in the innate response (Cao, 2016).

Once TF activity was estimated, the network structure prior was then used to bias model selection of TF-to-gene target regulatory interactions towards edges with prior information during network inference (Figure 3D). To improve the overall performance and stability of network inference, several aspects of the method (and key inputs) were tested, such as: different sources of priors (publicly available ENCODE data vs our ATAC-seq data), different TF binding motif databases (HOCOMOCO (Kulakovskiy et al., 2013) vs CisBp 2.0 (Weirauch et al., 2014)), and different model selection methods (Bayesian Best Subset Regression (BBSR) vs Elastic Net (EN)). Additionally, since every computational run is subject to stochasticity in the inference procedure, we evaluated whether network performance could be improved by combining hundreds of individual computation runs (selecting model components such as TF activity estimates that were stable across random subsamples of the structure prior; see STAR Methods). To evaluate the performance of networks inferred using the parameters described above, we used area under precision-recall curves (Madar et al., 2010) to compare the prediction and ranking of our TF-to-gene target regulatory edges against TRRUSTv2, a gold-standard reference database of human transcriptional regulatory events (Han et al., 2018), and well-known lists of ISGs (Kane et al., 2016; Schoggins et al., 2011; Shaw et al., 2017). We found that the EN regression model outperformed BBSR when ATAC-seq based networks were benchmarked against TRRUSTv2 (Figures 3E-F). We further improved precision-recall on TRRUST and known ISGs by bootstrapping 400 individual EN-ATAC inference runs into a converged network (EN-ATAC x400) (Figures 3E-H; S3C-D). This final EN-ATAC network correctly predicted 90 out of 97 “core” mammalian ISGs (Shaw et al., 2017) to be downstream of IRF and NF-κB family members (Figure S3E) and emphasized a role for IRF3 (Figure S3F). Thus, we were able to integrate gene expression and chromatin accessibility data to estimate a network with 542 TFs and 21,862 target genes, explaining > 2/3 of the variance in our expression dataset (Table S5).

### Network clustering, differential gene expression analysis, and TF enrichment tests define key subregulatory groups

We have used this DC transcriptional response network to group the expression changes observed into sets of co-regulated genes with high confidence regulatory subnetworks. We first applied a cutoff to visualize the top 75k regulatory edges that displayed the highest confidence beta scores (Figure 4A) and then partitioned the network using Louvain-modularity clustering into 10 major “neighborhoods” of TF and target gene communities, with each cluster encompassing at least 1% of the total number of genes in the network (DC network topology can be freely explored using Gephi software by downloading the Gephi-formatted supplemental file, available in the STAR Methods section). We were able to assign putative biological functions to 7 out of 10 clusters based on pathway enrichment scores (see STAR Methods). The most striking enrichment scores were observed in the top right shoulder of the network, demarcating clusters that are predicted to function in the interferon response (Cluster 5, IRFs & Interferon), inflammation and cytokine production (Cluster 8, NF-κB & Inflammation), and in response to xenobiotic stress (Cluster 2, Regulation of Cell Activation) (Figure 4B-D).

**Figure 4.**
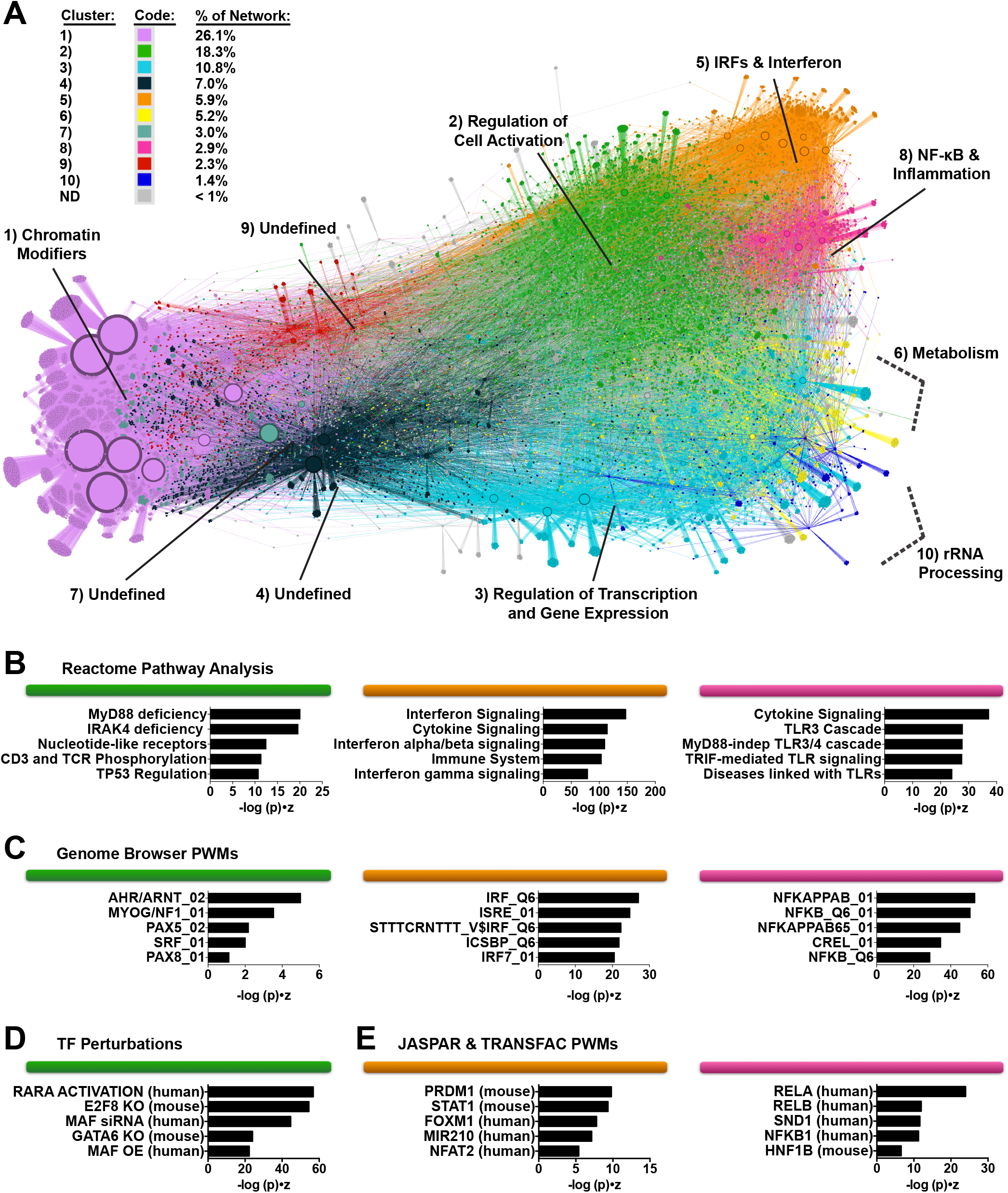
Modularity clustering of the network defines TF and gene target clusters predicted to control distinct biological functions. (A) EN-ATAC network topology in DCs visualized through Gephi software. Clusters were named according to pathway and motif enrichment (see STAR Methods). Arrows indicate transcriptional regulatory events between TFs and targets and do not specify positive or negative effects on target expression. Node size denotes relative number of edges. Clusters are color-coded based on Louvain clustering and ranked based on their decreasing size (from 1 to 10) (based on % of genes in each cluster vs total number of genes) (B-D) Enrichment of pathways from Reactome (B), Genome Browser TF position weight matrices (PWM) (C), TF Perturbations (D), and JASPAR and TRANSFAC PWMs (E) for Clusters 2, 5, and 8 (color-coded as in panel A). Ranking is by Enrichr combined score: (-log (p) * z-score). See also Figure S4 and Tables S5-6.

Activation of IRF, STAT, and NF-κB family members is centrally linked to interferon signaling during the innate immune response (Cao, 2016; Ivashkiv and Donlin, 2014). Fittingly, we found that several IRF (IRFs 1, 2, 3, 5, 7, 8, and 9), STAT1 & STAT2, and all five NF-κB (RELA, RELB, REL, NFKB1, and NFKB2) populated the upper right shoulder of the network (Figure S4A), consistent with their overlap in motif preferences (Figure S4B). Interestingly, IRF4 and IRF5, which also have GAA- and GAAA-rich motif features similar to that of other IRFs (Figure S4C), are found positioned in different areas of the network. IRF4, in particular, is localized to the extreme lower left (Figure S4A; Cluster 1, Chromatin Modifiers) in proximity to pioneer factors and repressors that target large numbers of genes (Jankowski et al., 2016).

We next sought to establish the relative impact of each TF in the network during an innate response. Towards this end we performed hypergeometric tests to assess TF enrichment, and found high enrichment for IRF, STAT, and NF-κB family members during stimulation with HIV-1-GFP, LPS, and pIC (Figure S4D; Table S6), with kinetics closely matching what was observed for IFN production and DC maturation (Figures S1A, S1D). We also found mild enrichment of these TFs during LKO-GFP infection, supporting the concept that non-replicating lentiviral vectors are partially, if inefficiently, sensed by the innate immune system (Figure S4D) (Johnson et al., 2018). By highlighting network topology where genes are differentially expressed during treatment with HIV-1-GFP, LKO-GFP, LPS, and pIC, we determined that the majority of the transcriptional response is concentrated in Clusters 2, 5, and 8 (Regulation of Cell Activation, IRFs & Interferon, and NF-κB & Inflammation) (Figure S4E).

### Exploration and validation of modulators of the interferon response

The first wave of the antiviral response is driven by production of type I and type III IFNs (Levy et al., 2011). In DCs the major type I and type III IFNs that are expressed are *IFNB1* and *IFNL1*, respectively (Figure S1D). To visualize all high-confidence edges for IFNB1, IFNL1, and their regulators that were predicted by the network, we generated a subnetwork visualization tool (Figure 5A; see STAR Methods for instructions on how to access the Jupyter subnetwork widget). Upstream of both *IFNB1* and *IFNL1*, we identified IRF3 as well as additional IRFs that have been reported to contribute to their induction under various conditions (such as IRF1, IRF5, IRF7, and IRF8) (Ivashkiv and Donlin, 2014)). Several unexpected factors were also predicted to be upstream regulators of *IFNB1* and *IFNL1* (Figure 5A), and of these TFs, we note that HIVEP1, CBFB, STAT2, PRDM1, and KLF13 were expressed at relatively high levels in DCs and in some cases exhibited dynamic expression changes during the innate response (Figure 5B).

**Figure 5.**
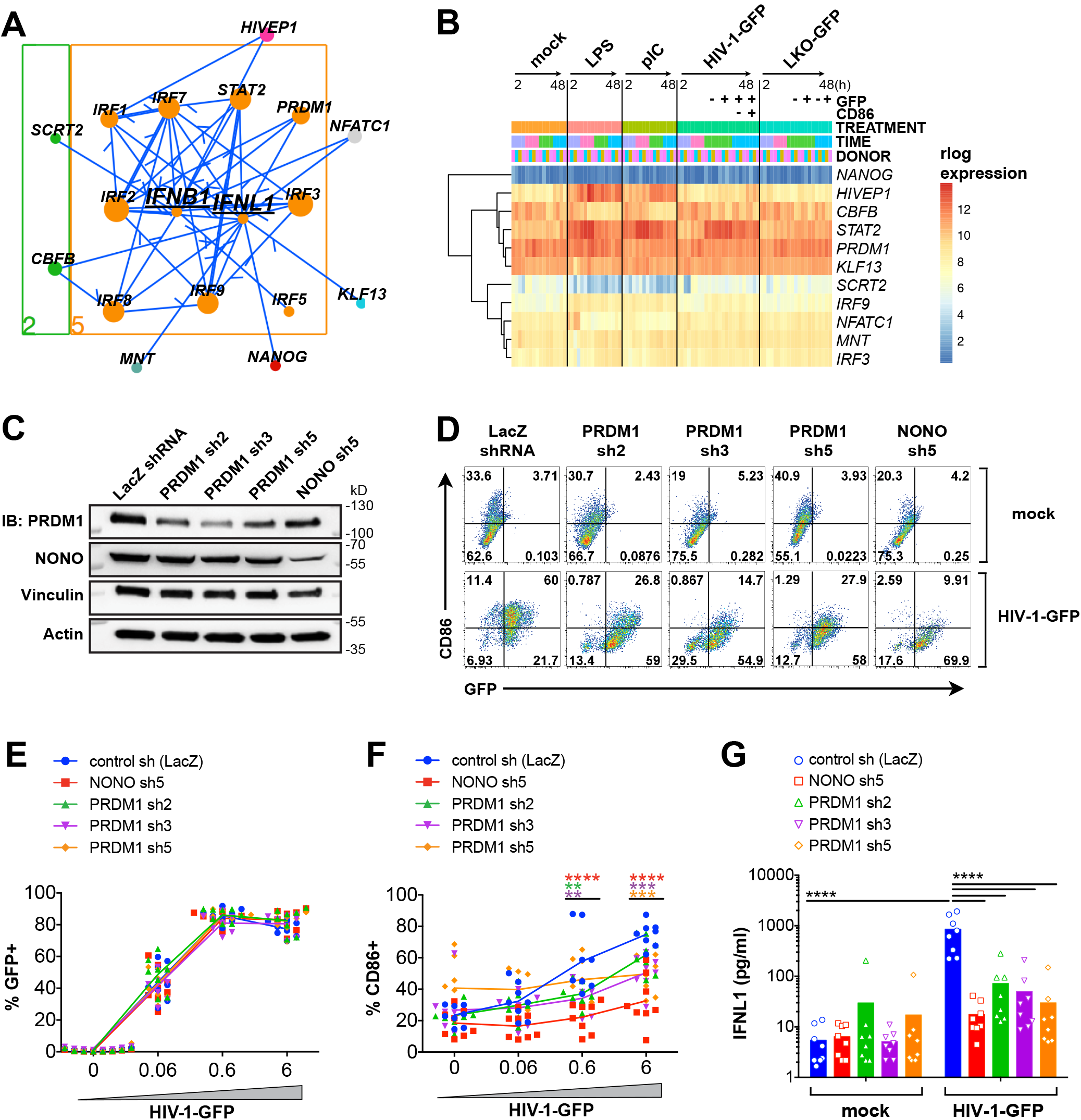
Subnetwork exploration uncovers positive and negative regulators of IFN and ISGs. (A) Jupyter widget subnetwork view (see STAR Methods) of predicted upstream regulators of *IFNB1* and *IFNL1.* Arrows indicate transcriptional regulatory events between TFs and targets and do not specify positive or negative effects on target expression. Node size denotes relative number of edges. Nodes are color coded by network cluster (see Figure 4). (B) Heat map of gene expression for the indicated TFs during treatment with LPS, pIC, HIV-1-GFP, and LKO-GFP at 2, 8, 24, and 48 h. (C) Immunoblots of DC lysates after treatment with the indicated shRNAs. (D) Flow cytometry of CD86 vs GFP expression in shRNA-modified DCs 48 h after mock treatment or infection with HIV-1-GFP. (E-F) Plots showing %GFP+ (E) and %CD86+ (F) from DCs treated as in (D). (G) ELISA of IFNL1 expression in supernatants from shRNA-modified DCs that were mock treated or infected with HIV-1-GFP for 48 h. For E-G, plots represent pooled data from 8 donors. ** p < 0.01; *** p < 0.001; **** p < 0.0001. See also Figure S5.

We next sought to empirically test the function of several of these TFs in DCs, noting that PRDM1 and HIVEP1 were among the highest ranking TFs according to hypergeometric tests (Figure S4D). Using shRNA vectors we targeted these TFs for knockdown in DCs along side key sensors of HIV (cGAS and NONO) and the essential TF, IRF3. Among other potential TF candidates, we additionally tested KLF13, based on its relatively high expression and known role in recruiting coactivators p300/CBP and PCAF to drive expression of CCL5 (Ahn et al., 2007). We confirmed knockdown of these targets either by immunoblot ((Figure 5C) or qPCR (Figure S5A), and as expected, knockdown of cGAS and NONO potently inhibited DC maturation during HIV-1-GFP infection (Johnson et al., 2018; Lahaye et al., 2018; Lahaye et al., 2013) (Figures 5D-F; S5B-E). The most effective shRNA clones for HIVEP1, KLF13, and PRDM1 had no effect on viral infection. Under these conditions, we could not conclude whether HIVEP1 knockdown impacted DC maturation (Figure S5D). Knockdown of KLF13 partially inhibited DC maturation, which was significant for two shRNAs at high HIV-1-GFP MOI (Figure S5E). Moreover, we found that greater KLF13 knockdown correlated with lower levels of CD86 and inhibition of *IFNL1* expression during HIV-1-GFP infection (Figure S5F). In a more dramatic way, knockdown of PRDM1 significantly inhibited CD86 induction and IFNL1, and these effects were observed with three independent shRNAs (Figure 5F-G). Together, these data suggest that while cellular factors such as cGAS, NONO, and IRF3 function as critical toggle switches during the innate response, additional TFs such as KLF13, and, to a greater extent, PRDM1, may operate to fine-tune the magnitude of the innate response. Furthermore, other TFs with high enrichment scores identified from experiments with wild type HIV-1, HIV-2, recombinant interferon, and discrete innate immune stimuli (Figure S5G; Table S6) warrant further exploration.

### Mature and immature DC populations cannot be distinguished based on IRF/STAT enrichment alone

IRF and STAT activation is critical for inducing an interferon gene signature, but neutralization of IFN signaling does not completely block upregulation of inflammatory cell markers during HIV infection (Lahaye et al., 2013; Manel et al., 2010), suggesting that other factors in addition to these TFs contribute to DC maturation. Gene set enrichment analysis on CD86-low and CD86-high sorted DCs (Figures S1A; S2B) identified “DC Maturation” genes as significantly enriched in CD86-high compared to CD86-low populations (Figure 6A-C). However, we found no statistically significant difference in “Hallmark IFN Response” genes, with ISG expression levels and chromatin accessibility elevated in both conditions. (Figure 6B and S6A-B). Thus, an IFN response is not sufficient to explain the differences in DC maturation status following HIV infection.

**Figure 6.**
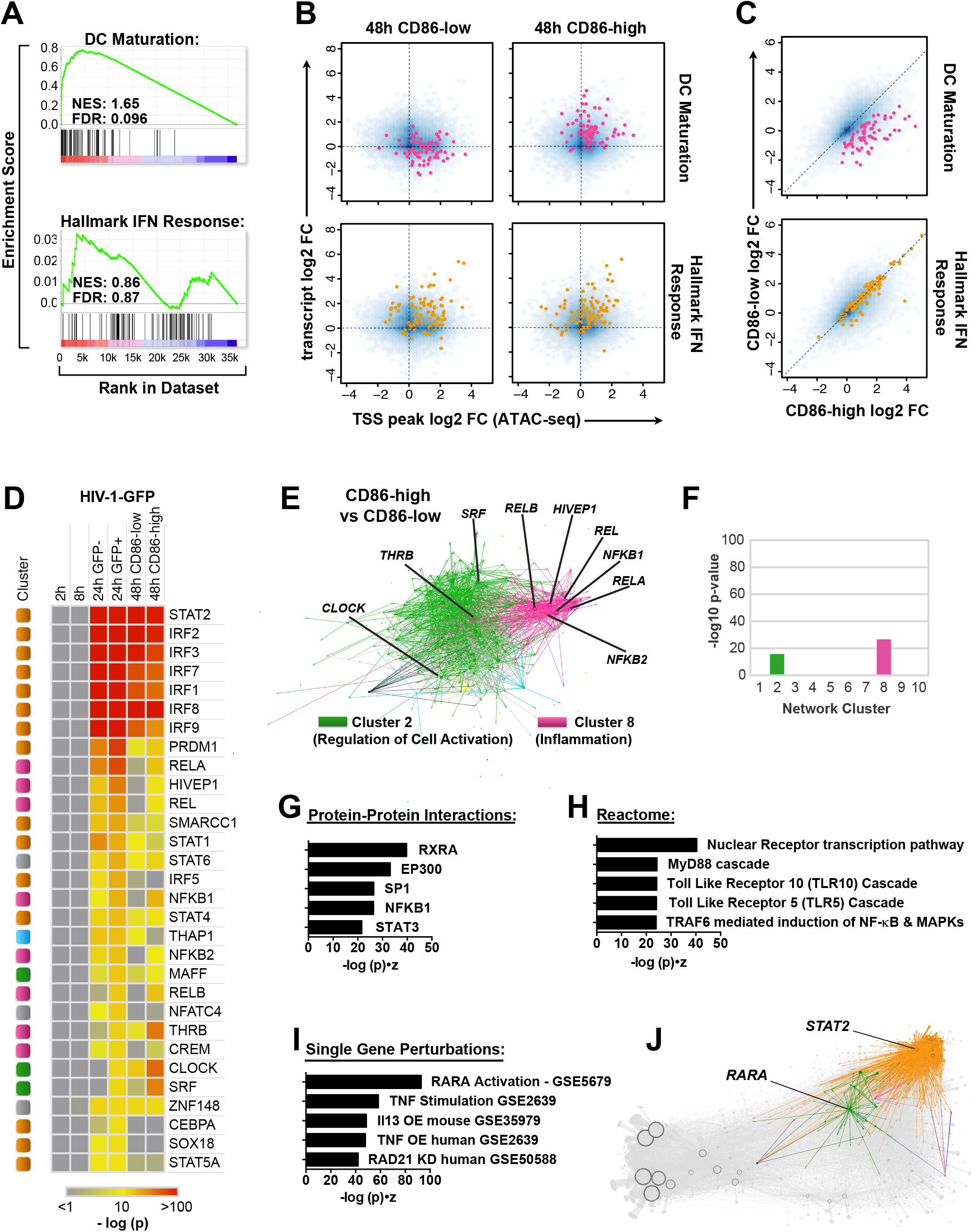
Mature DCs cannot be identified based on an IFN signature alone. (A) GSEA plots comparing gene expression data for CD86-high (mature) vs CD86-low (immature) DCs. Enrichment is considered significant at FDR < 0.25 (B) Smooth scatter density plots of transcript levels (RNA-seq) vs. TSS chromatin accessibility (ATAC-seq) in HIV-1-GFP infected DCs for CD86-low and CD86-high populations. Blue density marks the entire genome. Genes in the DC Maturation gene set (pink) and the Hallmark IFN Response gene set (orange) are highlighted in the top and bottom plots, respectively. (C) Log2 normalized gene expression plotted for the CD86-low population (y-axis) and CD86-high population sorted at 48 h. DC Maturation genes (pink) and Hallmark IFN Response genes (orange) are shown as in (B). (D) Heat map of hypergeometric scores for TF enrichment displaying HIV-1-GFP compared to mock at 2 h, 8 h, 24 h (GFP-), 24 h (GFP+), 48 h (CD86-low), and 48 h (CD86-high). TFs were ranked in descending order according to the 24 h GFP+ condition. Heat map is color coded according to –log p-value of TF enrichment. (E) Differential gene expression contrast between CD86-high and CD86-low populations as visualized by Gephi software, highlighting edges in Clusters 2 and 8. Ranking is by Enrichr combined score: (-log (p) * z-score). (F) Enrichment of differentially expressed genes separated by network cluster for the CD86-high vs CD86-low contrast as in (E). Potential Protein-Protein interactions (G), Reactome pathway enrichment (H), and Single Gene Perturbations (I) predicted from differentially expressed genes in the CD86-high vs CD86-low contrast. (J) Network connections for RARA and STAT2 visualized by Gephi software, color coded by contrast, and superimposed onto a grey network backdrop. See also Figure S6.

Consistent with our “DC Maturation” gene set results, the NF-κB family members (RELB, REL, NFKB1, RELA, and NFKB2) showed stronger enrichment in CD86-high than in CD86-low populations (Figure 6D-F) as did additional, unexpected, TFs (CLOCK, THRB, SRF, and HIVEP1) found in clusters 2 (Regulation of Cell Activation) and 8 (NF-κB & Inflammation). Interestingly, pathway analysis of differentially accessible chromatin in CD86-high vs CD86-low conditions suggested that Retinoic Acid Receptor Alpha (RARA), PPARD, NCOR1, and the canonical NF-κB family member RELA (among other TFs) influenced the transition between immature and mature DCs (Figure S6C-E). When differentially expressed genes were assessed in CD86-high vs CD86-low conditions in similar fashion, pathway analysis predicted strong enrichment of protein-protein interactions with the RARA partner molecule, RXRA, pathways associated with nuclear receptor transcription, MyD88 signaling, and gene expression changes resembling published datasets from RARA perturbation (highest among other inflammatory activators) (Figures 6G-I). These analyses implicated RARA/RXRA nuclear hormone receptor signaling to be involved in regulating DC maturation.

#### Identification of RARA as a critical TF for controlling DC maturation

At steady-state, RARs bind retinoic acid response elements (RAREs) together with RXRs in a complex with nuclear corepressors (NCOR1/2) and suppress transcription of RARE-dependent genes (Cunningham and Duester, 2015). Ligand binding leads to the dissociation of NCOR1/2 from RAR receptors and the recruitment of coactivators, which then drive gene expression. Given that RARA occupies a central region of Cluster 2 (Regulation of Cell Activation) (Figures 4D and 6J) and was predicted to influence DC maturation, we sought to assess its network connections and function compared to STAT2 and cGAS. Subnetwork visualization of RARA and close network edges displayed many connections between Cluster 2 (Regulation of Cell Activation) and Clusters 5 and 8 (IRF/Interferon and NF-κB/Inflammation, respectively) (Figure 7A). Many, but not all of these connections were detected using the Search Tool for the Retrieval of Interacting Genes (STRING) database (Szklarczyk et al., 2015), highlighting the position of RARA in close proximity to TFs involved in controlling cell fate, lipid & sterol metabolism, interferon signaling, and inflammation (Figure 7B).

**Figure 7.**
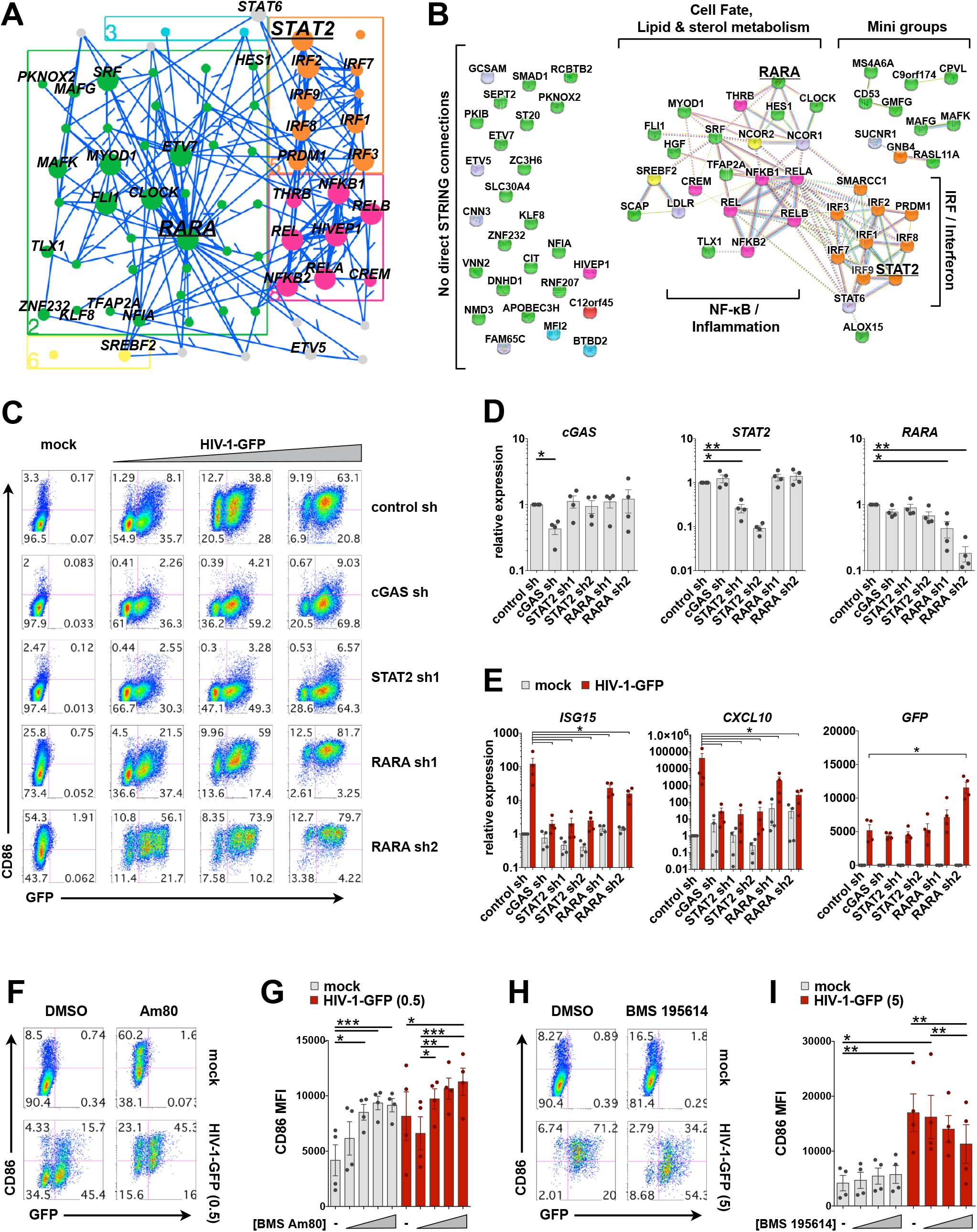
RARA is a negative regulator of DC maturation. (A) Jupyter widget subnetwork view (see STAR Methods) showing edges between RARA, its predicted targets, STAT2, and the remaining top 30 TFs ranked by hypergeometric p-value. Arrows point from TFs to targets but do not suggest positive or negative activity. (B) STRING database connections for nodes shown in (A) including NCOR1 and NCOR2. Brackets indicate groups and biological pathways. Nodes connected in the EN-ATAC network with no direct STRING connections are shown to the left. In (A) and (B) nodes are color coded by network cluster. (C) Flow cytometry plots of DCs modified by shRNA that were mock infected or challenged with HIV-GFP at day 4 for 48 h. Plots show CD86 vs GFP expression and represent 1 of 4 donors (MOI = 0.5, 1.5, 5). (D) qPCR validation of target knockdown in DCs modified by the indicated shRNAs. (E) qPCR of *ISG15, CXCL10*, and *GFP* expression in shRNA-modified DCs that were either mock treated or infected with HIV-1-GFP (MOI = 5) for 24 h. (F) Flow cytometry of DCs treated with vehicle or the RARA agonist Am80 (10 µM) ± infection with HIV-1-GFP for 48 h. (G) Pooled data of CD86 MFI in DCs from 4 donors treated as in (F). Am80 concentrations = 10nM, 100nM, 1 µM, 10 µM. (H) Flow cytometry of DCs treated with vehicle or the RARA antagonist BMS 195614 (10 µM) ± infection with HIV-1-GFP for 48 h. (I) Pooled data of CD86 MFI in DCs from 4 donors treated as in (H). BMS 195614 concentrations = 1nM, 100nM, 10 µM. For D, E, G, and I: n = 4 donors. * p < 0.05; ** p < 0.01; *** p < 0.001. Data represent mean ± SEM. See also Figure S7.

Knockdown of cGAS or STAT2 led to a pronounced loss of DC maturation during infection with HIV-1-GFP (Figure 7C-D). In contrast, knockdown of RARA upregulated CD86 at baseline and potentiated DC maturation during infection, suggesting RARA acts as a negative regulator of inflammation. Consistent with this notion, expression levels of many predicted targets of RARA increased in response to LPS and pIC treatment and to a lesser degree in response to HIV-1-GFP (Figure S7A). Moreover, we found an inverse relationship between gene expression of *RARA* and costimulatory molecules *CD80* and *CD86* (Figure S7B), supported by the negative correlation between the estimated transcriptional activity of RARA and its measured gene expression (Figure S7C). Knockdown of RARA was associated with a slight but statistically significant decrease in production of interferon and interferon stimulated genes *ISG15* and *CXCL10* (Figure 7E and S7E-H), leading us to consider that loss of RARA might impact the balance between antiviral interferon responses and inflammatory responses that benefit virus replication. In agreement with this idea, we observed heightened virus transcription in RARA knockdown conditions (measured by GFP reporter expression, Figures 7C; 7E), as increased inflammation is a correlate of increased HIV replication (Deeks et al., 2013).

Direct modulation of RARA using the pharmacological agonist Am80 (Delescluse et al., 1991) triggered increased CD86 surface expression both at baseline and following HIV-1-GFP infection (Figure 7F-G). We also observed that treatment with Am80 or the classic RARA ligand, all-*trans* retinoic acid (ATRA), increased gene expression of *CD86* (Figure S7D). Conversely, use of the RARA antagonist BMS 195614 reduced upregulation of CD86 during virus infection (Figure 7H-I). Although our findings from RARA agonists and antagonists may appear inconsistent with that of RARA knockdown, they are in line with the dual functions of nuclear receptors in recruiting either coactivators or corepressors of transcription, which will be further discussed below. Together, these data support our network predictions that spotlight a role for RARA in the regulation of DC maturation.

## Discussion

Multiple transcription factors (TFs) are known to drive the innate response downstream of pathogen sensing in myeloid DCs (Ivashkiv and Donlin, 2014), but it remains unclear how production of interferon, differential expression of thousands of host genes, and DC maturation are coordinately regulated across the genome in response to HIV detection. To address this issue, we have employed a systems biology approach to understand how DCs respond to stimulation, generating steady state and time series RNA-seq under a battery of viral and innate immune perturbations combined with ATAC-seq measurements of chromatin accessibility (Figure 1A). We inferred a transcriptomic network linking 542 TFs to 21,862 targets and report 500k predicted regulatory interactions. To date, this represents one of the largest gene regulatory networks that has been charted in mammalian cells.

Computational strategies that combine ATAC-seq and gene expression data have recently been employed to identify key regulators of tissue-resident memory CD8+ T cells (Milner et al., 2017), define TF specificity across cell types (Cusanovich et al., 2018), and chart the landscape of human cancers (Corces et al., 2018). Here, we improved the stability of network inference by performing ensembles of analyses, with each run subsampling information from a structured network “prior” that was generated from our ATAC-seq data. By modeling changes in gene expression to be a function of combined TF activities, we reduced the complexity of multiple factors known to influence gene expression into a single parameter (such as changes in TF expression, post-translational modifications, and epigenetic marks on target genes). In doing so, our network revealed strong enrichment of TFs known to regulate the innate response, notably including IRF3, a key factor that is activated by phosphorylation and, as it is constitutively expressed, is not often identified in pathway analyses that emphasize changes in TF expression (Amit et al., 2009).

Our final gene regulatory network exhibited a high level of precision/recall against TFs targeting known ISGs and correctly predicted prominent roles for IRF-, STAT-, and NF-κB family members in the IFN response. We recovered 90 out of 97 “core” mammalian ISGs in the top 75k network edges as targets of IRFs, and note that the remaining 7 core ISGs were predicted to be downstream of STAT- and AP-1 family TFs (Table S5). Interestingly, of the 22 different IFN species measured by RNA-seq, we found *IFNL1* to be upregulated more than any type I IFN, including *IFNB1,* during stimulation with LPS, pIC, or HIV-1. We know IFN signaling can provide both beneficial and detrimental effects during progression to AIDS (Sandler et al., 2014), yet the contribution of type-I vs type-III IFN in an *in vivo* response to HIV remains to be determined.

Among several TFs predicted to be upstream of IFN genes in our network, we found that PRDM1 (also known as Positive Regulatory Domain-I, PR/SET Domain-1, and BLIMP1) is an essential positive regulator of innate immune activation in response to HIV-1 infection in DCs. For almost 30 years, PRDM1 has been known to bind IFN promoter sequences (Keller and Maniatis, 1991). Since then, reports have suggested that PRDM1 is important for differentiation of plasma cells, is a risk factor for autoimmune disease, and can act to either positively or negatively regulate cell-specific cytokine production (Bonelt et al., 2019; Ko et al., 2018; Nutt et al., 2007). Here, we have revealed that PRDM1 promotes IFNL1 expression and DC maturation during HIV infection, in accordance with its high hypergeometric enrichment and cluster association with key IRFs in the network. Future exploration of our network could help uncover whether factors associated with PRDM1 determine how PRDM1 operates to promote or inhibit IFN production in a cell-specific manner.

By analyzing RNA-sequencing data from sorted cell populations, we have determined that induction of a robust ISG signature is insufficient for DC maturation, suggesting that factors in addition to PRDM1, IRF-, and STAT-family TFs members influence the transition from immature to mature DCs (Figure 6). In support of this notion, we have demonstrated that the retinoic acid receptor, RARA, acts as a negative regulator of DC maturation at basal state (Figures 7 & S7). Based on our network analysis predicting that RARA and NF-κB activities govern the transition from immature to mature DC, together with our data indicating that RARA modulates innate immune responses, we propose a model in which, at basal state, RXR/RARA heterodimers bind to RAREs and inhibit DC maturation driven by NF-κB family members (Figure S7I). Perturbation of RARA expression, either by shRNA or unknown factors during the innate response relieves this inhibition and permits DC maturation (Figure S7J). Binding of ligand/retinoids to RARA results in the exchange of bound nuclear corepressors with nuclear coactivators that will also permit DC maturation. Thus, RARA knockdown and administration of RARA agonists can both achieve a similar biological end, albeit through different means (Figure S7K). In agreement with this model, using an RARA antagonist to block the release of nuclear corepressors inhibits DC maturation in response to HIV infection (Figure S7L). These results reinforce the reported involvement of retinoids in DC maturation (Geissmann et al., 2003; Szatmari and Nagy, 2008). Given that retinoic acid is known to activate MAPK phosphorylation and abrogate tolerance in a stressed intestinal environment (DePaolo et al., 2011), it is not unreasonable to envision that RARA activation might influence derepression of inflammatory gene expression during HIV infection.

Our model thus suggests that RARA function could provide another layer of control to govern DC maturation through sensing changes in metabolism. Differentiation of DCs *in vitro* and *in vivo* is known to depend on fatty acid synthesis (Rehman et al., 2013). In addition, in plasmacytoid DCs, lipid metabolism, RXR, and PPAR-related pathways can modulate type I IFN responses, suggesting that environmental cues such as retinoids help shape immune responses (Wu et al., 2016). It is tempting to draw parallels with how changes in fatty acid oxidation and cholesterol biosynthesis influence innate responses and determine cell fate (tolerogenic versus immunogenic DC) (Maldonado and von Andrian, 2010; York et al., 2015).

In addition to profiling how DCs respond to HIV, it is noteworthy to mention that we have profiled responses to a battery of classic innate immune stimuli, including LPS, pIC, cGAMP, R848, and recombinant interferon. This represents a valuable source of information that one can explore to compare similarities and differences among transcriptomic pathways bridging innate immune detection and production of inflammatory cytokines. We note that much of the transcriptomic activity observed for HIV infection in our network overlaps with that in response to LPS and pIC exposure; however, we found that HIV drives unique changes in fatty acid oxidation pathways (Johnson et al., 2018). These findings support the existence of an interplay between HIV infection, lipid metabolism, and retinoids that control IFN production and DC maturation, which will require further investigation.

Additionally, we note that HIV-1 infection drives transient changes in chromatin accessibility that likely precede or perhaps coincide with the induction and resolution phases of the innate response. Interestingly, we observed global changes in chromatin accessibility that were not always associated with changes in host gene expression. We speculate that this could reflect on either the nature of the innate response or on signals derived from events in the virus life cycle, as many of these changes were located in gene-dense regions known to be hot-spots for virus integration (Table S3, (Wang et al., 2007)).

Collective analyses of these data will help to further decipher the mechanisms of innate immune regulation that span initiation, feed-forward amplification, and the resolution of aberrant hyperactivation. Understanding DC maturation and IFN production will likely inform future antiviral strategies, since the robust activation of T cells, which is facilitated by mature DCs, is crucial for immunological control of HIV (Walker and McMichael, 2012). We anticipate that the quality of network inference will further improve as motif databases are refined and large-scale genetic perturbations become tractable in primary human DCs. The network presented herein highlights how the innate response depends on the coordinated action of multiple TFs and serves as a resource for further exploration of pathways that govern HIV replication and innate immunity.

## Supporting information

Table_S2_Expression_matrix_all_samples

Table_S3_ATAC_peak_RLOG_counts

Table_S4_Prior_matrix

Table_S5_EN_ATAC_Network_500k_edges

Table_S6_Hypergeometric_tests_all_contrasts

## Acknowledgments

This project was supported the Howard Hughes Medical Institute (DRL); the Helen and Martin Kimmel Center for Biology and Medicine (DRL); by grants from the National Institutes of Health (F32AI093231 to JSJ; R21AI084633 to DRL; R01AI025032, R01AI032972, and U19AI100627 to AA); State funding from the Agence Nationale de la Recherche under “Investissements d’avenir” program (ANR-10-IAHU-01) (MMM) and ATIP-Avenir INSERM program (MMM).

## Author Contributions

Conceptualization, JSJ, AR, NM, DRL, RB, and MMM; Methodology, JSJ, ND, AR, and MMM; Software, ND, AR, AW; Formal Analysis, ND, AR, and LA; Investigation, JSJ, XL, SL, and MMM; Writing – Original Draft, JSJ and MMM; Writing – Review & Editing, all authors; Resources, AA, NM, DRL, RB and MMM; Supervision, NM, AA, NM, DRL, RB, & MMM; Funding Acquisition, JSJ, AA, NM, DRL, RB, and MMM.

## STAR Methods

### CONTACT FOR REAGENT AND RESOURCE SHARING

Further information and requests for resources and reagents should be directed to and will be fulfilled by the Lead Contact, Mickaël M. Ménager.

### EXPERIMENTAL MODEL AND SUBJECT DETAILS

#### Cell lines and blood-derived dendritic cells

To generate monocyte-derived immature DCs, we acquired leukocytes from de-identified normal human donors from a variety of sources (Bloodworks Northwest, Renton, WA, USA; New York Blood Center, New York, NY, USA; ARUP Blood Services, Sandy, UT, USA; and from venipunctures (approved by the Institut National de la Santé et de la Recherche Médicale ethics committee, Paris, France). The authors cannot report on the sex, gender, or age of the donors since the samples were de-identified and donors remain anonymous. Peripheral blood mononuclear cells (PBMCs) were layered over Ficoll-Paque Plus (GE Healthcare). CD14+ monocytes from PBMC buffy coats were isolated with anti-human CD14 magnetic beads (Miltenyi) and cultured in RPMI (Thermo Fisher) containing 10% heat-inactivated fetal bovine serum (FBS, Peak Serum, Inc), 50 U/ml penicillin, 50 µg/ml streptomycin (P/S, Thermo Fisher), 10 mM HEPES (Sigma), 2-Mercaptoethanol (Thermo Fisher), and 2 mM L-glutamine (Thermo Fisher), in the presence of recombinant human GM-CSF at 10 ng/ml and IL-4 at 50 ng/ml (Peprotech). Multiple lots of FBS were tested to identify batches leading to minimal baseline induction of CD86 over the course of DC differentiation. Fresh media and cytokines were added to cells (40% by volume) one day after CD14+ cell isolation. On day 4, cells were collected, resuspended in fresh media with cytokines used for infection or stimulation. Immature DCs on day 6 were routinely assessed by flow cytometry surface marker staining to be CD11c+ (Thermo Fisher Cat# 17-0116-42, RRID:AB_1659668), HLA-DR+ (BioLegend Cat# 307607, RRID:AB_314685), DC-SIGN+ (R and D Systems Cat# FAB161P, RRID:AB_357064), and CD86- (eBioscience Cat# 15-0869-42 RRID:AB_11042003). DC experiments were performed using biological replicates from blood-derived cells from multiple individual donors as indicated in the figure legends.

293FT female cells (Life Technologies Cat# R70007, RRID:CVCL_6911) were cultured in Dulbecco’s modified Eagle’s medium (DMEM, Thermo Fisher) that was supplemented with 10% FBS, P/S, 10mM HEPES, and supplemented with 0.1 mM MEM non-essential amino acids (Thermo Fisher), 6 mM glutamine, and 1 mM sodium pyruvate (Thermo Fisher).

HL116 male cells were cultured in DMEM supplemented with 10% FBS, P/S, 10mM HEPES, 0.1 mM MEM non-essential amino acids, 6 mM glutamine, 1 mM sodium pyruvate, and HAT supplement (Thermo Fisher; hypoxanthine (5 mM), aminopterin (20 µM) and thymidine (0.8 mM)) diluted 1:50 as recommended.

All cell lines were thawed from early passages, kept in culture no longer than 4 weeks, and were regularly tested for mycoplasma contamination (every 6 months). All cells were maintained at 37°C and 5% CO2.

### METHOD DETAILS

#### Plasmids

HIV-1-GFP has been used previously to study immune responses in human DCs (Manel et al., 2010) and is env- vpu- vpr- vif- nef-, with the GFP open reading frame in place of nef. We generated virus like particles packaging Vpx from the plasmid pSIV3+ (based on SIVmac251, GenBank acc. no. M19499), which has been described elsewhere (Mangeot et al., 2000).

Target sequences for shRNA vectors are listed in Table S7. All lentiviral constructs were transformed into Stbl3 bacteria (ThermoFisher) for propagation of plasmid DNA. All plasmids were prepped and purified using Nucleobond Xtra Maxi Kit (Takara) or similar maxi prep reagents.

#### Virus and virus-like particle production

To produce recombinant lentiviral vectors, reporter viruses, and Vpx-containing virus-like particles, we transfected plasmids into 293FT cells as previously described (Johnson et al., 2018). Briefly, the day before transfection, cells were seeded onto poly-L-lysine (MP Biomedicals) -coated 15 cm plates to be 70-80% at the time of transfection. Cells were transfected with a total of 22.5 µg DNA using PEImax (Polysciences, Inc.) at a ratio of 1:2 (DNA:PEI). For lentiviral vectors, plasmid amounts were 3.4 µg CMV-VSV-G (Addgene plasmid #8454), 9 µg psPax2 (Addgene plasmid #12260), and 10.1 µg transgene (LKO.1 controls or shRNA construct). For HIV reporter viruses plasmid amounts were 3.4 µg CMV-VSV-G and 19.1 µg HIV cassette. Virus-like particles containing Vpx were produced using 3.4 µg CMV-VSV-G and 19.1 µg pSIV3+ (Mangeot et al., 2000). The morning after transfection, cells were washed once and fed with 30 ml of fresh media. Supernatants were harvested 32-36 h after feeding 293FTs and then passed through 0.45 µm syringe filters (Corning) to remove debris. Supernatants of reporter virus and lentiviral vectors were either used fresh for transduction or concentrated by ultracentrifugation by spinning in conical tubes (Beckman) at 25 K rpm for 2 h at 4 °C in an SW32 swing-bucket rotor (Beckman). Lentiviral pellets were thoroughly resuspended (∼30-fold concentrated) in DC media without cytokines. Before transducing cells, insoluble material was clarified from lentivirus stocks by centrifuging at 300 rcf for 3 min at 4C. Viral stocks were frozen at −80 °C and used as indicated below.

#### Perturbation of DCs

To genetically modify DCs, we isolated CD14+ monocytes from buffy coats as described above and transduced them with lentiviral vectors in the presence of virus-like particles packaging Vpx similar to previously described protocols (Johnson et al., 2018; Manel et al., 2010). Briefly, monocytes were resuspended in complete DC media containing GM-CSF (10 ng/ml), IL-4 (50 ng/ml), polybrene (Sigma, 1 µg/ml), and supernatants containing Vpx particles (1 ml supernatant for 1 × 10^7 cells) and then were plated in either 10 cm dishes (6 × 10^6 cells in 9 ml media) or 96 well plates (160,000 cells in 150 µl media). Roughly 30 min after plating, clarified concentrated lentiviral stocks were added to transduce cells. We routinely observed that 95-99% of cells could be transduced using 150 µl concentrated vector in 10 cm plates or 5 µl concentrated vector per well in 96 well plates.

#### Infections and stimulations

DCs were infected or stimulated with innate agonists between day 4 and day 6 after differentiation. DCs were counted on day 4 and resuspended at 800,000 cell per ml in fresh medium with GM-CSF, IL-4, and polybrene (1 µg/ml) and then reseeded into appropriate culture vessels. For most assays, DCs were infected at a density of 800,000 cells per ml by diluting virus in DC media (without cytokines or polybrene) to a final volume normalized to controls. Innate and inflammatory stimuli were used at the indicated concentrations: polyI:C (InvivoGen, 10 µg/ml); LPS (List Biological Laboratories, INC, 1 ng/ml).

#### Flow cytometry

Infected or stimulated DCs were washed with phosphate buffered saline (PBS, Corning) and then exposed to LIVE/DEAD violet (ThermoFisher) in PBS for 15 min at 4 °C in the dark. Cells were either simultaneously stained for surface markers or first washed with PBS and then stained in FACS buffer containing 1% Bovine Serum Albumin (BSA, Roche) and 1 mM EDTA in PBS for 15-30 min in the dark at 4 °C. Cells were then washed with PBS and fixed with 0.5% paraformaldehyde (Electron Microscopy Sciences) diluted in PBS. Cells were either sorted on a FACSAria II or data was acquired on an LSR II flow cytometer (BD Biosciences). Data were then analyzed using FlowJo software (FlowJo LLC).

#### RNA-seq

DCs from multiple donors were infected or stimulated as indicated in the figure legends, stained with anti-human CD86 and a live-dead viability dye and then sorted into serum on a FACSAria II (BD Biosciences). Cells were lysed in TRIzol reagent (Thermo Fisher), RNA was isolated according the manufacturer’s instructions, and then samples were submitted to HudsonAlpha Institute for Biotechnology (https://hudsonalpha.org/) for library preparation and RNA-sequencing.

50bp-length single-end sequences were aligned to the human genome (hg38) using STAR version 2.4.2a. Samtools 0.1.19 was used to filter alignments to a MAPQ score threshold of 30. Counts per gene were called using featureCounts version 1.4.6 and gencode v24 genome annotation. Samtools 0.1.19 was used to filter alignments to a MAPQ score threshold of 30. We confirmed the presence of HIV-1 or HIV-2 sequences in infected samples yet these sequences were not taken into account during differential expression analysis. Differential expression analysis was performed separately for each RNA-seq experiment using DESeq2 version 1.10.1 and R version 3.2.3, using time and treatment together as the design parameter. Genes were considered differentially expressed in a given comparison if the FDR-adjusted p-value was below 0.1.

#### ATAC-seq

DCs from three unique donors were infected in the presence of Vpx to match infection conditions as performed for RNA-seq at 2, 8, 24, and 48 h after infection. Cells were stained for LIVE/DEAD violet and CD86 to assess activation status and sorted on a FACSAria II (BD Biosciences) according to the gating strategy established for RNA-seq. 50,000 sorted DCs from each condition were immediately prepped for ATAC-seq (Buenrostro et al., 2013). Cells were pelleted by spinning at 500 g for 5 min at 4 °C, resuspended in cold PBS, and then centrifuged again at 500 g for 5 min at 4 °C. Cell pellets were lysed in cold lysis buffer (10 mM Tris-HCl, pH 7.4, 10 mM NaCl, 3 mM MgCl2 and 0.1% IGEPAL CA-630) and immediately spun at 500 g for 10 min at 4 °C. The nuclear pellet was used directly for the transposition reaction by resuspending in a reaction mix (20 µL 2× TD buffer, 2 µL Tn5 transposase (Nextera DNA Library Prep Kit, Illumina) and 18 µL nuclease-free water). Transposition was performed at 37 °C for 30 min and then samples were immediately processed using MinElute kit (Qiagen) to isolate DNA.

To generate ATAC-seq libraries for sequencing, DNA fragments from MinElute samples were amplified by PCR using NEBNext PCR master mix (New England Biolabs) and custom Nextera primers (Buenrostro et al., 2013) (and Table S7) using the following conditions: 72 °C for 5 min; 98 °C for 30 s; and then cycling at 98 °C for 10 s, 63 °C for 30 s and 72 °C for 1 min. To stop amplification before saturation in order to reduce bias, we monitored the PCR reaction using SYBR Green (Thermo Fisher) during qPCR. Samples were cycled a total of 12-15 times. Libraries were sequenced on a NextSeq 500 (Illumina) using a 150 cycle high output kit. Unique sequence read pairs were aligned to the human genome (hg38) using bowtie2 (2.2.3), filtered based on mapping score (MAPQ > 30, Samtools (0.1.19)), and duplicates removed (Picard version 1.120). Only pairs that aligned uniquely and concordantly to non-mitochondrial human chromosomes were retained.

For establishing the network prior, a merged ATAC-seq Sam file was generated by combining all ATAC-seq conditions. Peak calling was performed with Peakdeck version 1.1 (parameters - bin 160, -STEP 25, -back 5000, -npBack 10000) using the start and end locations of the pairs to define fragment lengths, with a p-value of 1e-4 as the cutoff, outputting 87,681 unique, non-overlapping, high-confidence peaks. Putative binding events were discovered by finding motif occurrences using HOCOMOCO version 10 (Kulakovskiy et al., 2013) as the motif database, with position weight matrices for 601 of 640 human TFs discovered using FIMO (Memesuite version 4.10.1) with a p-value cutoff of 1e-4. When scored against our gold standards (see below) HOCOMOCO performed better than the curated CisBP 2.0 motif database (Weirauch et al., 2014) composed of JASPAR (Mathelier et al., 2016), TRANSFAC (Matys et al., 2006), and other motif collections, possibly due to low redundancy in motif position weight matrices. Bedtools (version 2.25) was used to generate a Markov model of order 1 from all peak intervals to use as a background peak file (parameter “—bgfile”). For each TF, the target set was further refined by filtering out binding sites below the top quartile, ranked by the FIMO p-value. TF binding sites were linked to a target gene if the peak fell within a 1kb window of the gene body, creating a prior matrix with 601 columns and 24909 rows (Table S4). Entries at row “I” column “J” were set to 1 to denote a putative regulatory event between TF “i” and target gene “j.” All other entries were set to 0, which penalizes, but does not completely prevent a regulatory edge connecting “i” and “j.”

#### Expression Normalization

Expression data was normalized using DESeq2 to remove the dependence of the variance on the mean. We explored linear models for batch correction such as ComBat and limma, but these models were ruled out as they superimpose each experiment’s center of mass – essentially placing different experiments on top of one another in PCA space—experiments which should have unique, non-overlapping centers of mass (due to having different ratios of stimulus conditions as compared to mock conditions). Thus, we used DESeq2 rlog-transformed expression data to normalized gene expression values relative to mock conditions for each time point and for each donor by doing a vector subtraction. In our time series RNA-seq experiment, 3 samples were identified as outliers in gene expression biclusters from DESeq2 analysis (noted in Metadata Table S1), consistently clustering away from the rest of the 93 samples across nearly all condition contrasts in PCA plots. Since network inference is sensitive to outliers due to a z-scoring step, and because follow-up RNA-seq experiments under the same conditions supported the idea that these samples were outliers, these 3 samples were removed from the network inference procedure.

#### Estimation of TF activities

The network inference methodology used here builds directly upon our previous platform, the Inferelator, in which the log gene expression is modeled as the linear combination of TF predictor variables (Bonneau et al., 2006). Earlier versions of the inferelator model used the TF expression as the predictor, but current methods use a latent variable, the estimated TF activity, as the predictor (Greenfield et al., 2013). The activity of a TF is inferred from the changes of the TF’s putative target genes and has been shown to be able to uncover more known relationships between TFs and target genes than using TF expression as the predictor (Arrieta-Ortiz et al., 2015).

In the Inferelator, we define the equation for inferring a network as X = BA. “X” is the gene expression matrix, “A” is the transcription factor activity matrix, and “B” is the betas matrix, which can also be thought as the connectivity network. If beta is found to be nonzero, then we have inferred an association between the gene and the TF. This linear equation appears straightforward, but it cannot be immediately solved because there are two unknowns, both “B,” the network beta value, and “A,” the TF activity. We therefore use a two-step process. First, we solve for “A,” using the prior network, with the equation X = PA. In this equation, the prior network “P” is known. However, because the confidence in any one of its chromatin accessibility derived entries is low, we randomly leave out half of the entries of P, and do this iteratively 400 times, deriving 400 estimates for the activities, “A.” We solve the inverse problem X = PA by multiplying “X” by the pseudoinverse of the prior matrix, “P”. The second step of the process is to solve the linear regression problem X = BA, with “B” as the unknown, which we do using stability selection with elastic net regularization.

#### Network inference with subsampled priors

Since the prior matrix was built with thresholding at fixed p-values, both at the level of peak finding and the level of motif binding, any single entry in the prior matrix cannot be considered accurate. In order to improve computational reproducibility, we use a subsampling strategy, which was recently pioneered in a different context to increase prediction accuracy when assessing whether pathway perturbations hold true across different animal species (Hafemeister et al., 2015). This subsampling strategy decreases the density of the prior matrix, which is a desideratum when the number of targets per TF in the prior is more than 2000 on average. Therefore, for each of the 400 computational estimations of the transcription factor activity, we sampled 50% of the nonzero entries in the prior without replacement. This enabled TFs with similar motif preferences and subtle differences in target sets to be teased apart over the course of many Inferelator runs. For example, in a single Inferelator run, a number of different IRF family members could be assigned at random to have undue emphasis on IFN and ISG expression, since many IRFs overlap in motif preference. In some extreme single run cases, due to inherent randomness in the inference procedure, no IRFs were assigned to IFNB1 or IFNL1. In contrast, after multiple runs where estimated TF activities were subsampled, the ATAC-seq-based network correctly predicted a strong influence of IRF3 on IFNB1, IFNL1, and core ISGs target sets were teased apart from other IRFs.

#### Ensemble Model and AUPR curves

The 400 individual models that were generated by the inferelator were combined by ranking each interaction by summed beta values to generate an ensemble network (EN-ATAC x400). While this ensemble model predicted over 2 million interactions, we have provided a table to accompany this manuscript that lists the top 500,000 edges, selected based on absolute value of the beta sign (Table S5). Networks of this size cannot be reasonably visualized or efficiently analyzed by current computational resources, so for this work we chose to display the top 75,000 edges in the ensemble network. At this choice of a cutoff in ranked interactions, there are 542 TFs assigned as regulators for 21862 targets. This ranking was used to compute an area under the precision-recall curve (AUPR) by validating the predicted interactions against “gold standard” TF-to-gene-target connections that have been reported in the literature.

#### Proportional Venn Diagrams

Venn diagrams were generated using BioVenn (Hulsen et al., 2008). IRFs 1, 3, 5, 7, 8, and 9 and the NF-κB family members RELA, RELB, REL, NFKB1, and NFKB2 are known to be important for primary and secondary IFN responses and influence ISG expression. We determined whether the targets of these IRFs and the NF-κB family members (Figure S3E), or specific targets of IRF3 that were predicted by the EN-ATAC x400 network or the BBSR network (Figure S3F) overlapped with a set of “core” mammalian ISGs (Shaw et al., 2017).

#### Hypergeometric tests for TF enrichment

With the network model established, we can then ask whether any given Transcription Factor is associated with differential gene expression through its network targets. For every contrast, we used the hypergeometric test for enrichment to query whether the set of differentially expressed genes for a given contrast was enriched for the targets of any of the transcription factors. We used a Bonferroni correction, dividing by the number of transcription factors, when estimating these p-values. For finding a global ranking of enrichment across multiple differential gene expression contrasts, the weighted z-method for combining probabilities was used with equal weights following a previously defined method (Whitlock, 2005).

#### Modularity and similarity K-means clustering

In order to interpret the results of the network inference, which yielded a network with 75,000 edges, we asked whether there existed large-scale clusters in the networks. To this end we used the python-louvain clustering module, which implements the modularity clustering algorithm, maximizing an objective function that sums edges within communities while subtracting edges between communities (Blondel, 2008). Due to the stochasticity of this algorithm, where the modularity score and even the number of clusters varies with each run, we repeated the run until convergence, which occurred on the order of a thousand runs. This computational approach allowed us not only to create clusters for large-scale communities, but also build a dendogram of pairwise similarities between every gene in the network, where the pairwise entry corresponds to the number of times in the thousand runs that the two genes were clustered together. This dendogram allowed us to identify “unclustered” nodes in the network: genes that do not consistently cluster with any of the main groups. The dendogram was split into 500, which resulted in 10 major clusters, defined as a cluster with more than 1% of the total gene set. These clusters were then labeled with identifiers from Enrichr pathway analysis. For complete pathway analysis results for the top 10 clusters, please see:

Cluster 1 http://amp.pharm.mssm.edu/Enrichr/enrich?dataset=3dhb9

Cluster 2 http://amp.pharm.mssm.edu/Enrichr/enrich?dataset=3dhbc

Cluster 3 http://amp.pharm.mssm.edu/Enrichr/enrich?dataset=3dhbo

Cluster 4 http://amp.pharm.mssm.edu/Enrichr/enrich?dataset=3dhfq

Cluster 5 http://amp.pharm.mssm.edu/Enrichr/enrich?dataset=3dhgc

Cluster 6 http://amp.pharm.mssm.edu/Enrichr/enrich?dataset=3dhgi

Cluster 7 http://amp.pharm.mssm.edu/Enrichr/enrich?dataset=3dhgn

Cluster 8 http://amp.pharm.mssm.edu/Enrichr/enrich?dataset=3dhgp

Cluster 9 http://amp.pharm.mssm.edu/Enrichr/enrich?dataset=3dhgq

Cluster 10 http://amp.pharm.mssm.edu/Enrichr/enrich?dataset=3dhgr

#### Network Visualization

Network visualization software Gephi Version 0.9.1 (Bastian, 2009) was used to visualize the network using the Force Atlas2 layout algorithm (Jacomy et al., 2014). For visualizing smaller network components with louvain clustering, we used a jp_gene_viz visualization tool developed at the Simons Foundation. The Gephi-formatted network file is available upon request.

#### Gene Set Enrichment Analysis (GSEA)

GSEA was performed by comparing each condition in the time series RNA-seq data to the corresponding mock time point. 36883 gene features for each condition were ranked by the signal to noise metric of GSEA and the analysis was performed using the standard weighted enrichment statistic against 3815 human gene sets contained in the Molecular Signatures Database that included all (H) Hallmark gene sets, (C2) curated gene sets, and (C3) motif gene sets. In a first pass analysis, the normalized enrichment score (NES) was calculated using 500 gene set permutations and once relevant gene sets were identified a more stringent statistical analysis was performed using 1000 phenotype permutations. Full GSEA results can be accessed through the following link: https://www.dropbox.com/sh/2u4psikt2tyz4r0/AAB4Re3YKtOA5lqPEEAwfR7Aa?dl=0

#### Immunoblotting

1 million cells were lysed in 100 µL of RIPA buffer (50mM Tris HCl, 150mM NaCl, 0.1% SDS, 0.5% DOC, 1% NP-40, Protease inhibitor (Roche; 1187358001)). Lysis was performed on ice for 30 minutes. Lysates were cleared by centrifugation at 8000g for 8 minutes at 4°C, and 20 µl of Laemmli 6X (12% SDS, 30% Glycerol, 0.375M Tris-HCl pH6.8, 30% 2-mercaptoehtanol, 1% bromophenol blue) was added and samples were boiled at 95°C for 15min. Lysates were resolved on Criterion or 4%-20% Biorad precast SDS-PAGE gels and transferred on PVDF membrane. Membranes were saturated and proteins were blotted with antibodies (listed in Key resources table) in 5% non-fat dry milk, PBS, 0.1% Tween buffer. ECL signal was recorded on the ChemiDoc-XRS or ChemiDoc Touch Biorad Imager. Data was analyzed and quantified with the Image Lab software (Biorad).

#### Quantitative PCR

50,000 to 200,000 cells were lysed in TRIzol reagent (Thermo Fisher) and then RNA was isolated following the manufacturer’s instructions with minor modifications. In brief, we performed two sequential chloroform extractions and added Glycoblue (Thermo Fisher) as a carrier prior to precipitation with isopropanol. RNA pellets were washed in 75% ethanol and resuspended in 200 µl of DNase and RNase-free water. 500 µg of RNA was converted into cDNA using Superscript III (ThermoFisher). Quantitative PCR reactions were carried out using TaqMan primer probes (ABI) and TaqMan Fast Universal PCR Master Mix (ThermoFisher) in either a Lightcycler (Roche) or a CFX96 thermocycler (BioRad) in a volume of 10 µl according to the following cycling conditions: 50 °C for 2 min, 95 °C for 2 min, then 55 cycles each of 95 °C for 3 sec, to 60 °C for 30 sec, followed by 95 °C for 5 sec. A melting curve analysis was then performed going from 65 °C to 95 °C in 0.5 °C intervals every 5 sec. Data were plotted as expression relative to *GAPDH* × 1000.

#### IFNL1 Protein Quantification

IFNL1 protein concentrations were measured on supernatants from infected or treated DCs using a LEGENDplex Human Anti-Virus Response assay (BioLegend) according to the manufacturer’s protocol. Data were acquired on a BD FACSVerse (BD) and analyzed with LEGENDplex Software (BioLegend).

#### Bioassays for type I IFN

To quantify IFN activity from infected or stimulated cells we assayed supernatants with HL116 reported cells that contain firefly luciferase gene under control of the IFN-inducible 6-16 promoter (Uze et al., 1994). Supernatants from treated or untreated DCs were transferred to 20,000 HL116 cells in 96 well plates. After 7 h, HL116 cells were lysed in passive lysis buffer and subsequently scored for luciferase activity using a luciferin-based method (Promega). Relative light units were converted to units per ml of IFN using a standard curve that was generated from serial dilutions of recombinant human IFNa2a, with HL116 cells responding in a linear range between 2 and 200 U/ml of IFN.

### QUANTIFICATION AND STATISTICAL ANALYSIS

Statistical analyses incorporated into the computational implementation of network inference were performed as stated above in the STAR METHODS section. Otherwise, statistical tests were performed as indicated in the figure legends or using Prism 6.0 (GraphPad) to calculate either a two-way ANOVA with Sidak’s multiple comparisons test or a two-tailed t test using paired samples. The number of unique donors for DC experiments is also listed in the figure legends, representing the number of biological replicates performed for a given experiment. Data reflects pooled data from multiple experiments where indicated.

### DATA AND SOFTWARE AVAILABILITY

The RNA-seq data have been deposited in the Gene Expression Omnibus (GEO) database under ID code GSE125817. The ATAC-seq data have been deposited in the GEO database under ID code GSE125918. Both can be accessed from the GEO series code GSE125919.

### KEY RESOURCES TABLE

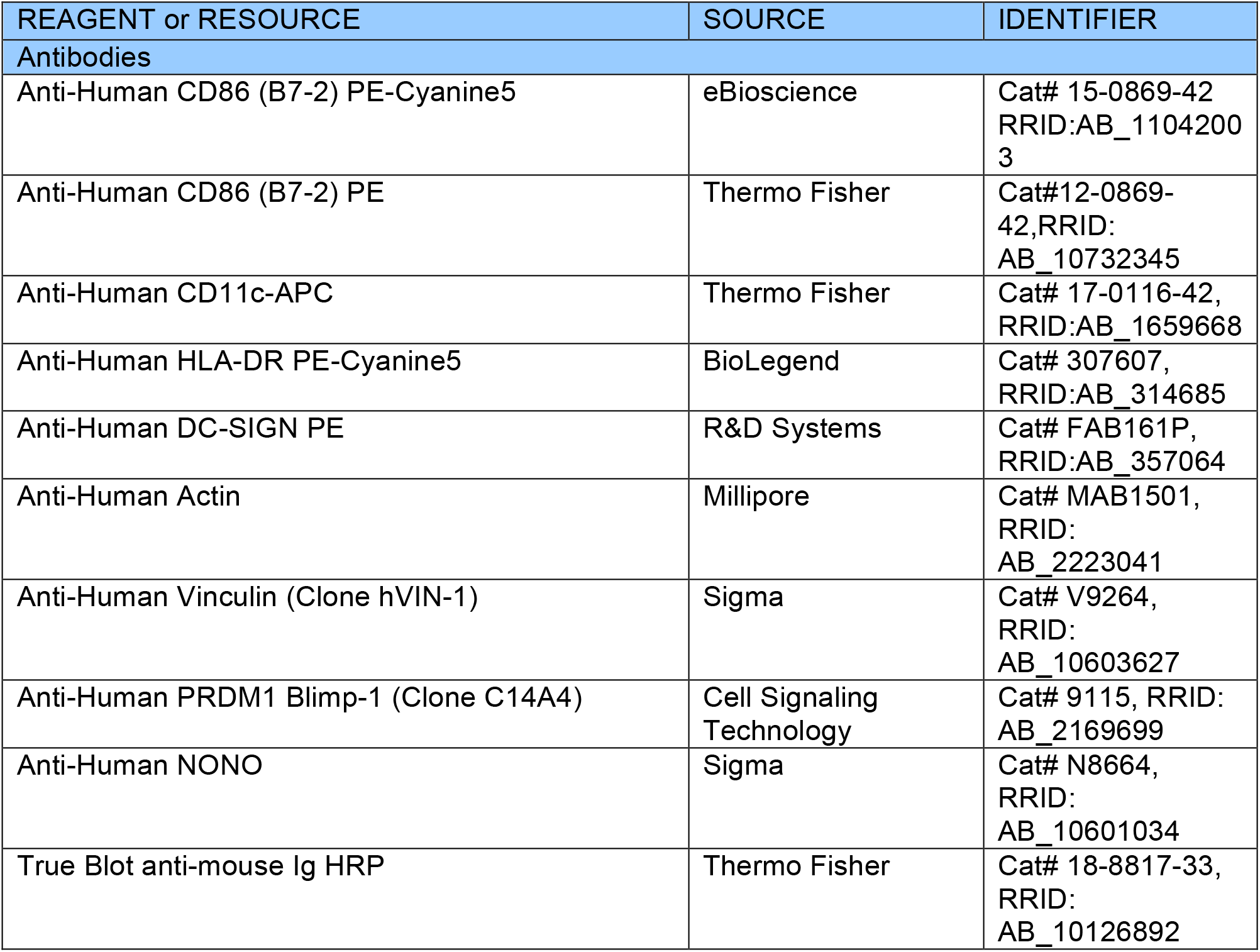
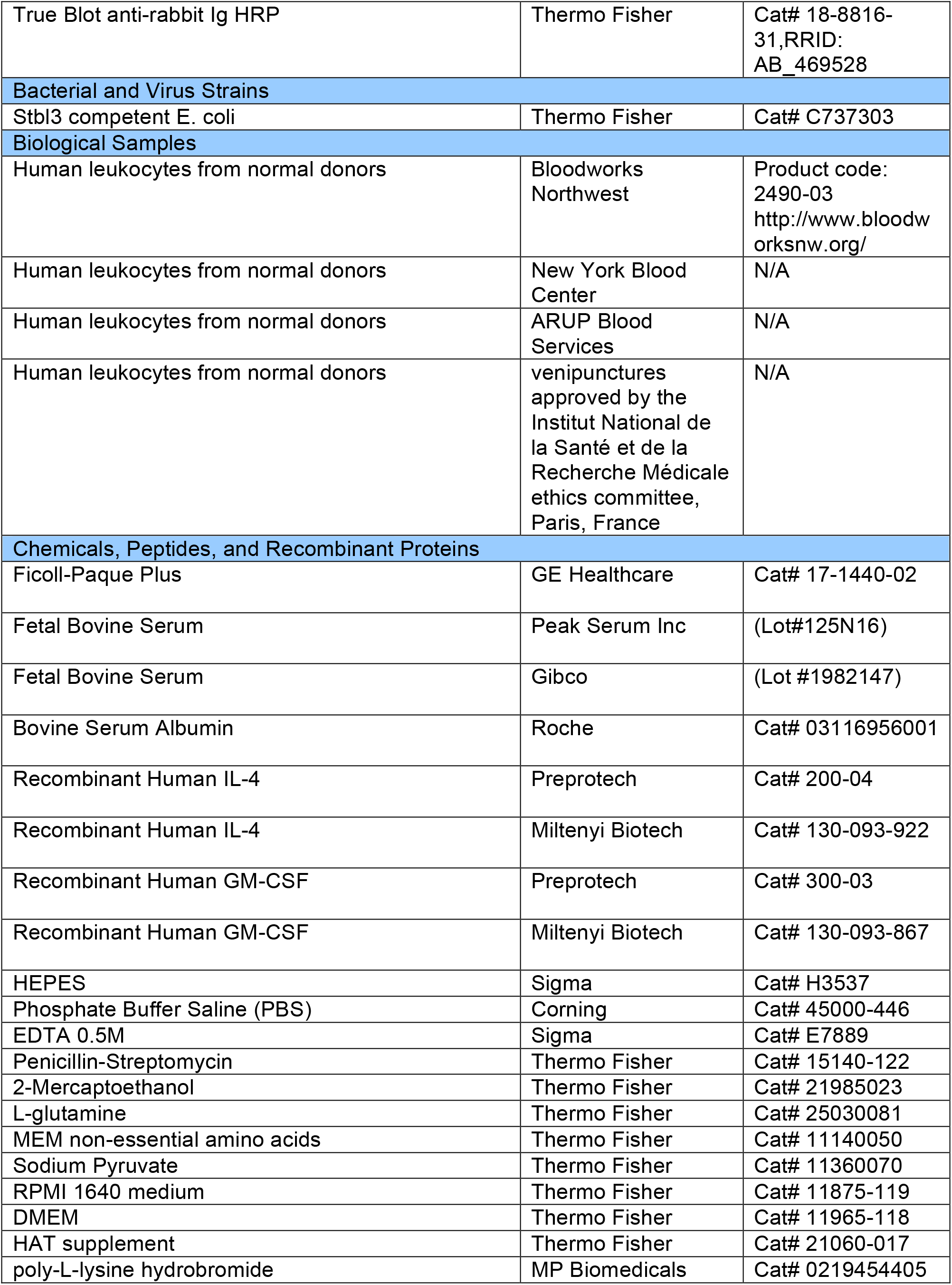
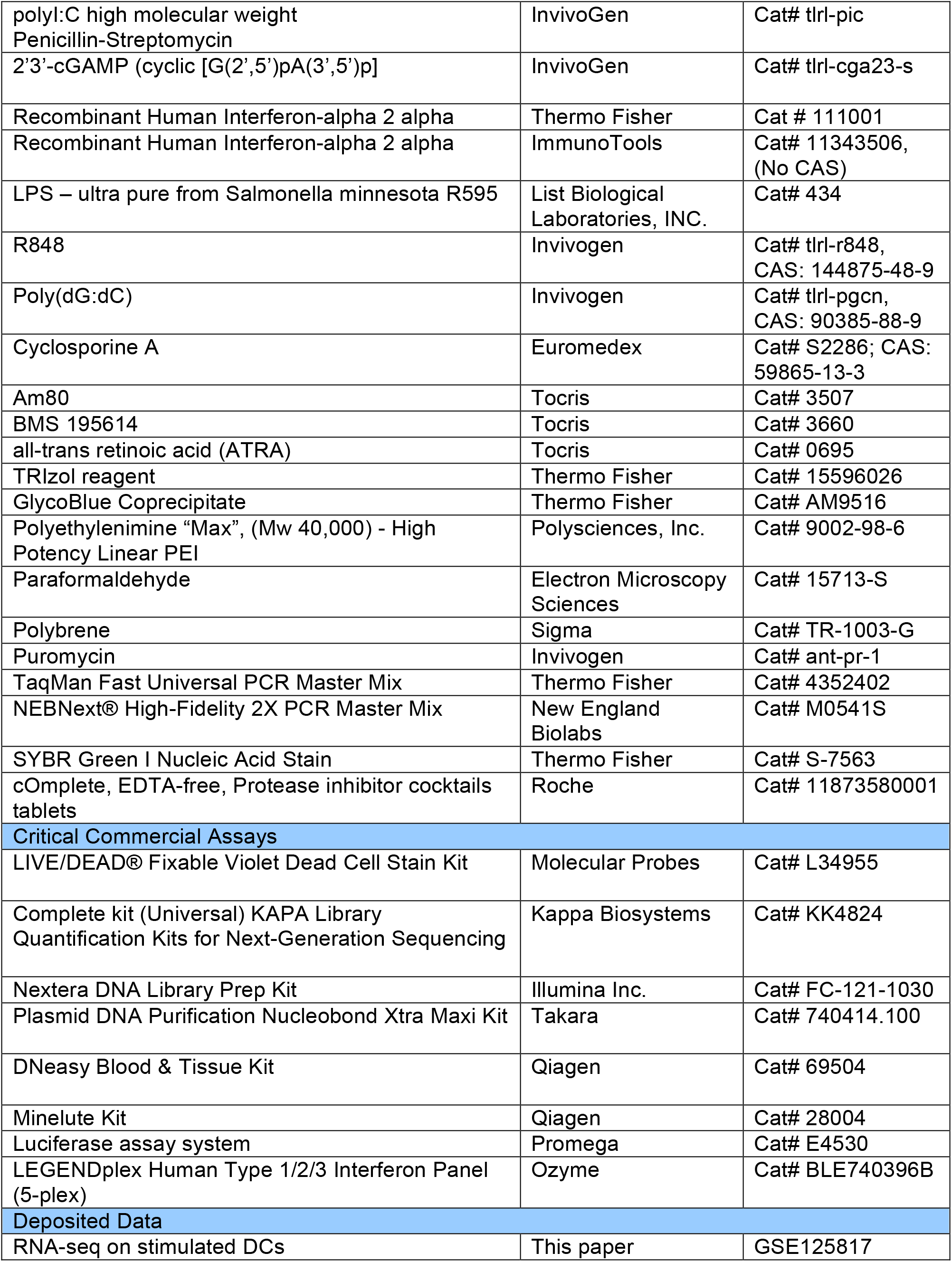
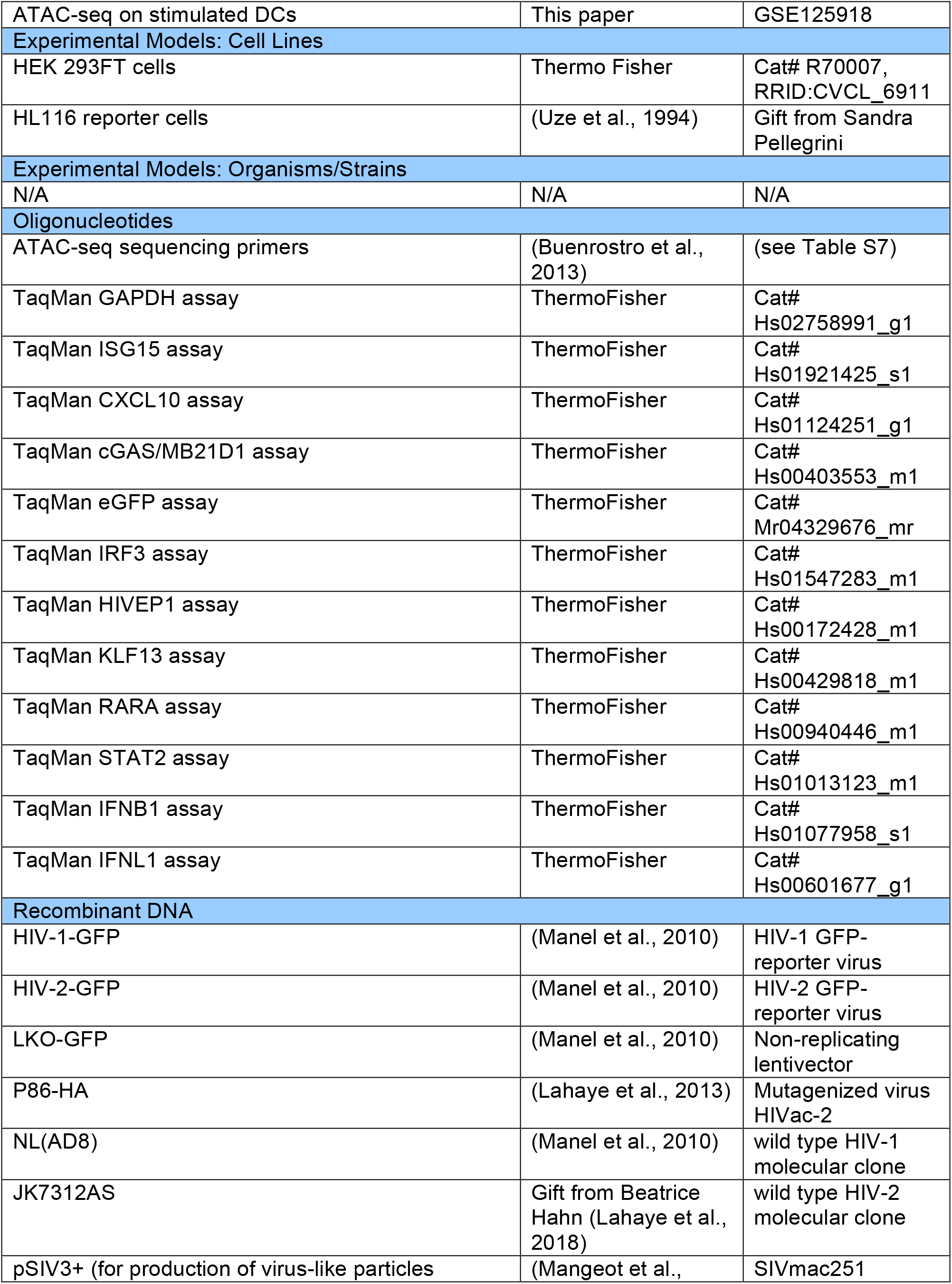
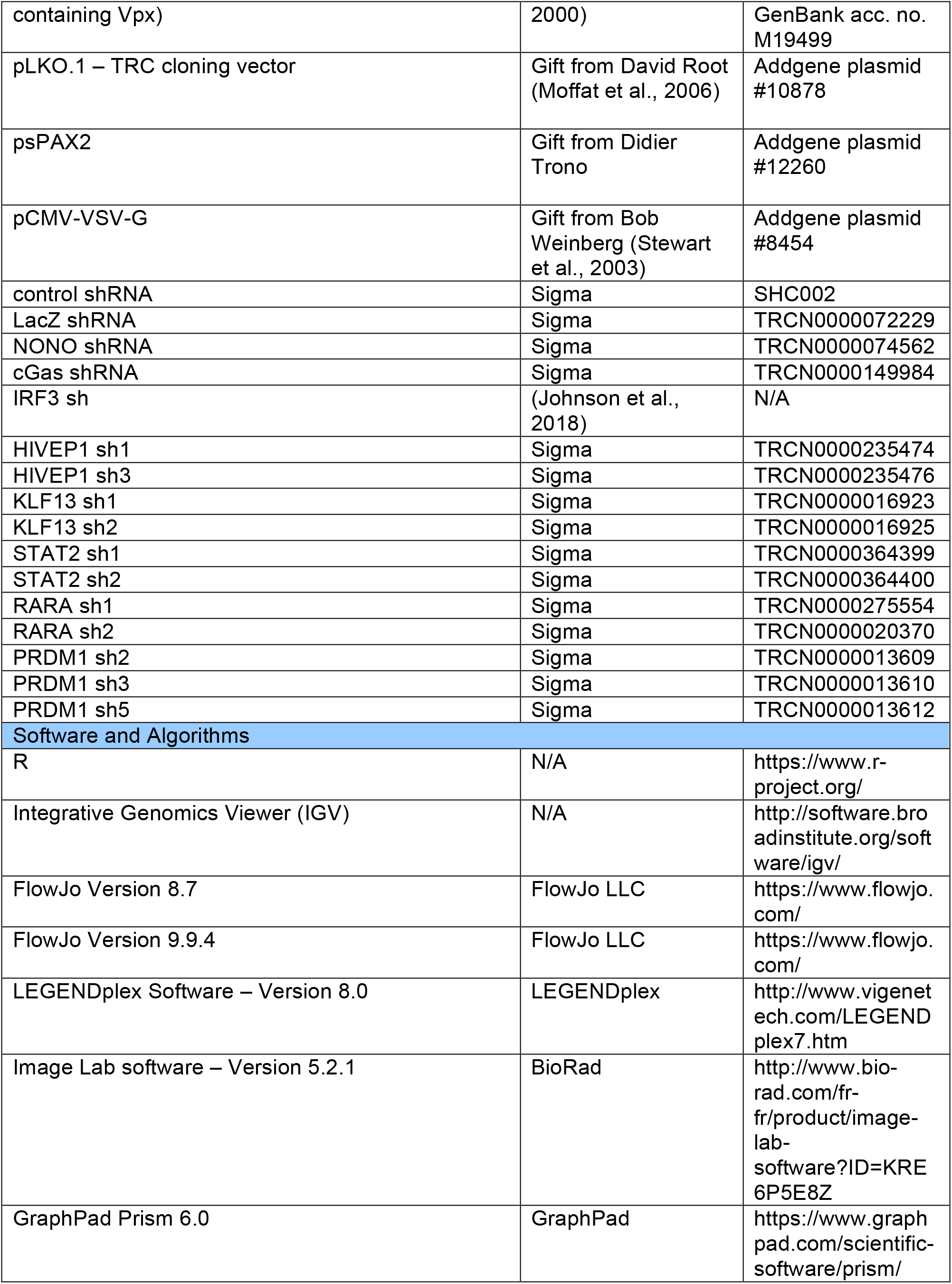
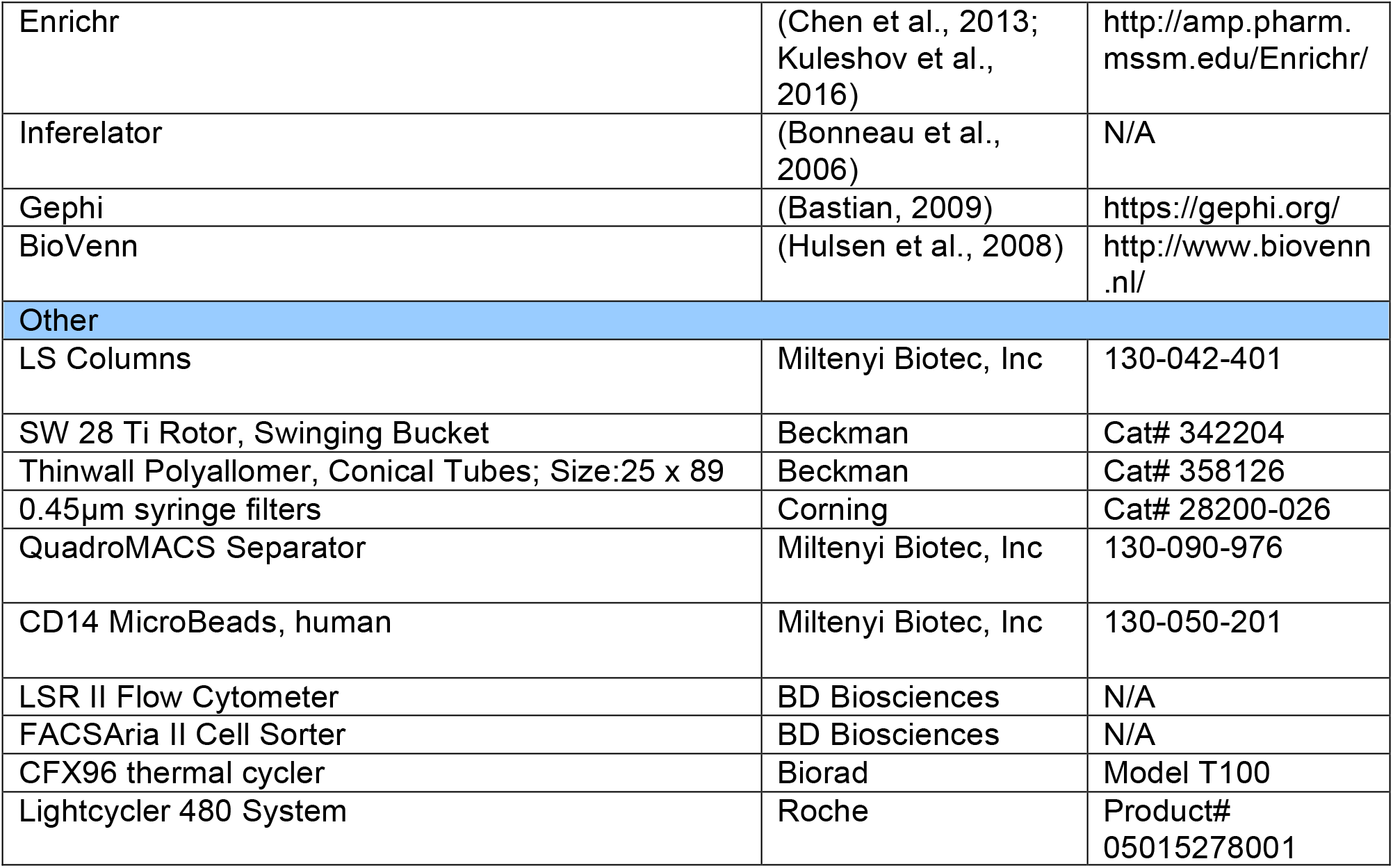

**Excel Tables not appended to the Supplemental PDF:**

**Table S2. Expression matrix. Related to Figure 1. Table S3. ATAC-seq peaks. Related to Figure 2. Table S4. Prior matrix. Related to Figure 3**.

**Table S5. DC gene regulatory network (EN-ATAC x400) capped at 500k edges. Related to Figure 4**.

**Table S6. TF Hypergeometric enrichment. Related to Figure 4**.

