## Supplemental information for "A comprehensive map of the dendritic cell transcriptional network engaged upon innate sensing of HIV"

Figure S1

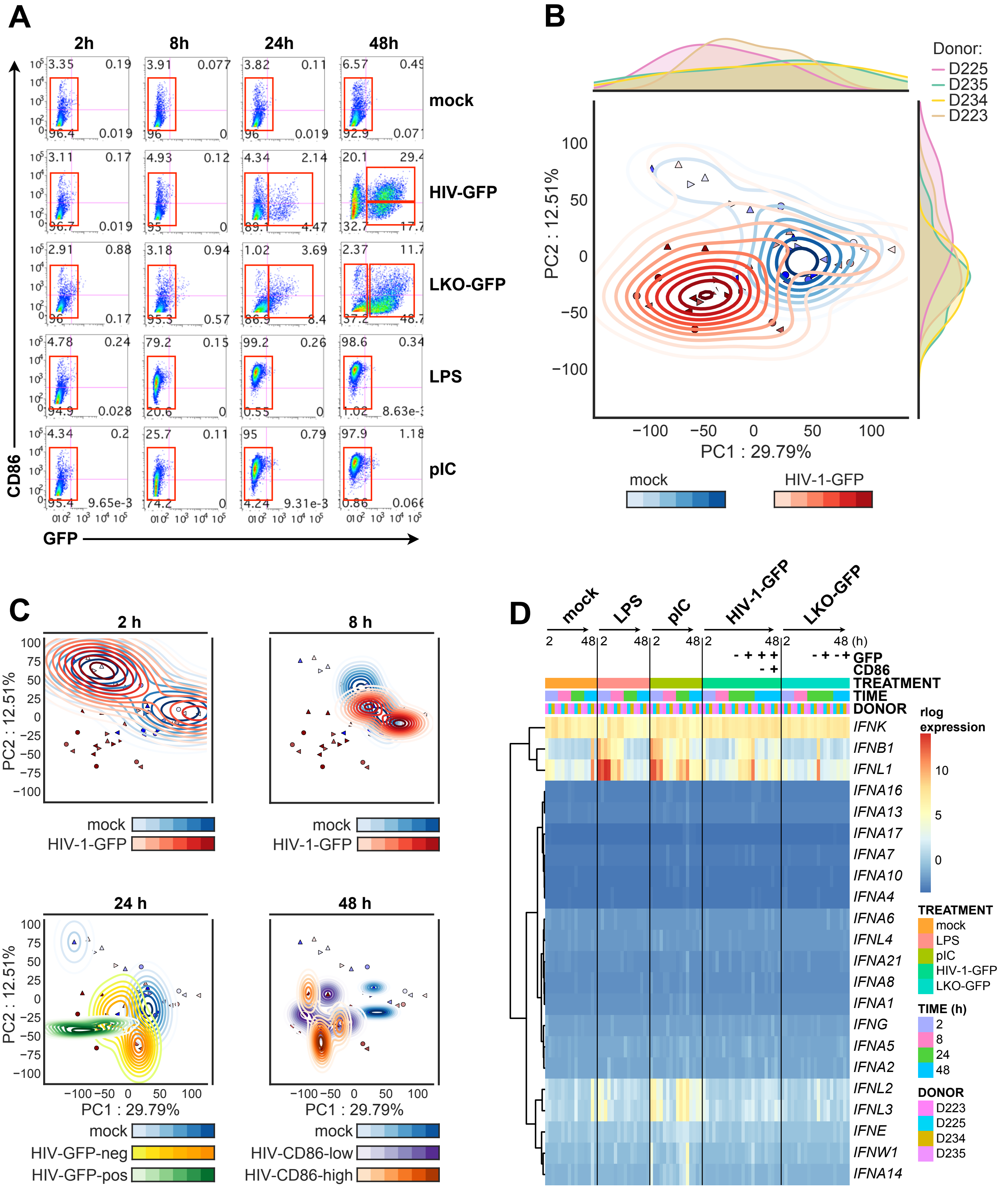

A
2h
8h
24h
48h
mock
HIV-GFP
LKO-GFP
LPS
pIC
CD86
GFP
B
Donor:
D225
D235
D234
D223
PC2 : 12.51%
PC1 : 29.79%
mock
HIV-1-GFP
C
2 h
8 h
24 h
48 h
mock
HIV-1-GFP
HIV-GFP-neg
HIV-GFP-pos
HIV-CD86-low
HIV-CD86-high
D
mock
LPS
pIC
HIV-1-GFP
LKO-GFP
GFP
CD86
TREATMENT
TIME
DONOR
IFNK
IFNB1
IFNL1
IFNA16
IFNA13
IFNA17
IFNA7
IFNA10
IFNA4
IFNA6
IFNL4
IFNA21
IFNA8
IFNA1
IFNG
IFNA5
IFNA2
IFNL2
IFNL3
IFNE
IFNW1
IFNA14
rlog expression
TREATMENT
mock
LPS
pIC
HIV-1-GFP
LKO-GFP
TIME (h)
2
8
24
48
DONOR
D223
D225
D234
D235

**Figure S1. IFN responses and DC maturation during HIV-1-GFP infection are delayed compared to LPS and pIC stimulation. Related to Figure 1.**

(A) Flow cytometry plots of DCs sorted for RNA-seq after infection with HIV-1-GFP at 2, 8, 24, and 48 h (MOI = 5). Plots show CD86 vs GFP expression and red boxes indicate sorted populations. Plots show representative data from 1 of 4 donors.

(B) PCA plots for RNA-seq data from DCs that were either mock treated or infected with HIV-1-GFP. All time points were grouped by condition (shades of blue or red for mock and HIV-1-GFP, respectively) showing mean-centered contour lines to mark standard deviations. Marginal curves show the density of sample position for each donor.

(C) PCA plots for RNA-seq samples as in (B) showing mean-centered contour lines for conditions separated by time (2, 8, 24, and 48 h). Color scale bars indicate mock, HIV-1-GFP, and specific populations.

(D) Heat map of expression data for 22 human IFN genes from time series RNA-seq samples depicted in (A).

Figure S2

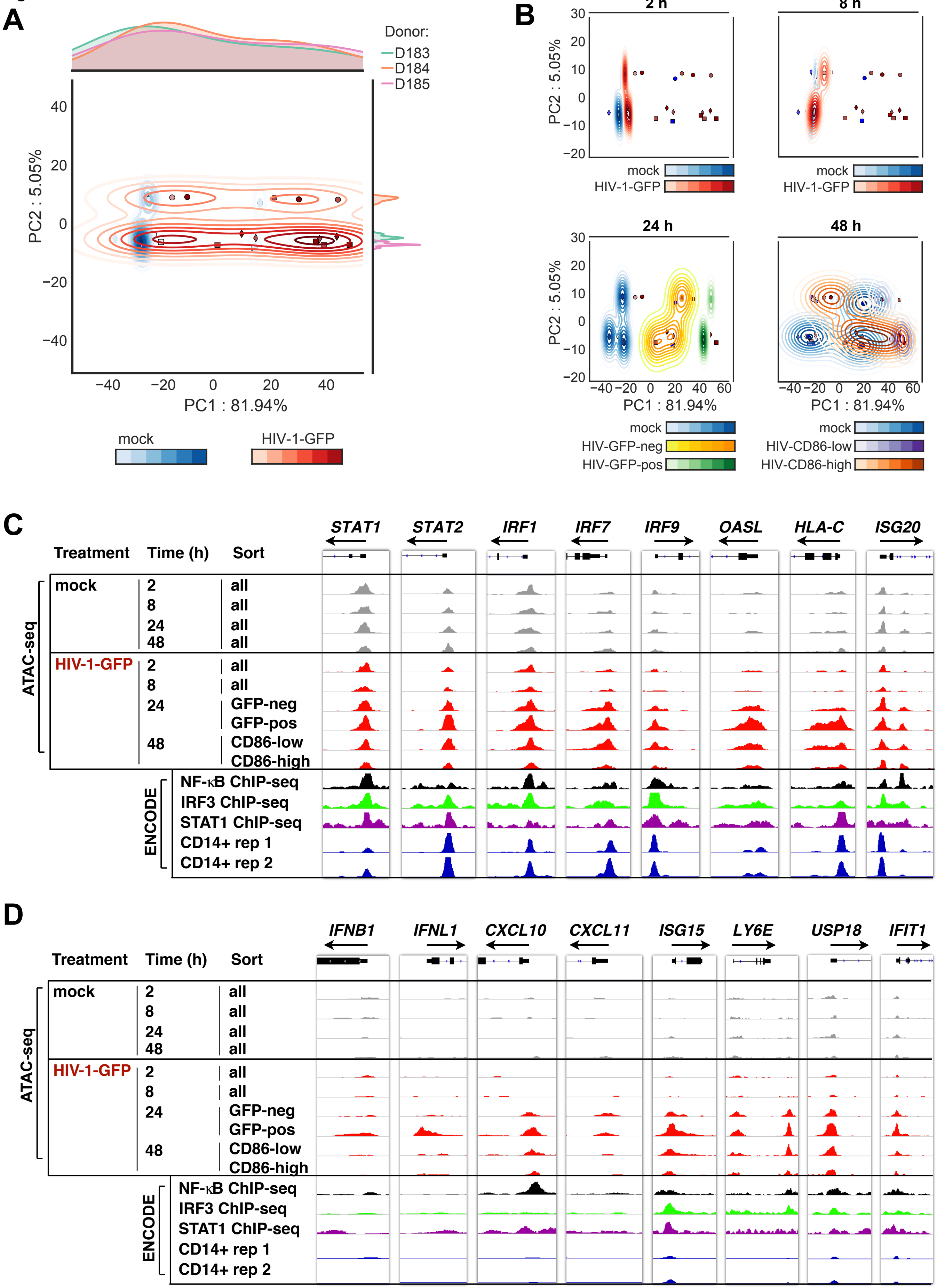
A
Donor:
D183
D184
D185
PC2 : 5.05%
PC1 : 81.94%
mock
HIV-1-GFP
B
2 h
8 h
24 h
48 h
mock
HIV-1-GFP
HIV-GFP-neg
HIV-GFP-pos
HIV-CD86-low
HIV-CD86-high
C
STAT1
STAT2
IRF1
IRF7
IRF9
OASL
HLA-C
ISG20
Treatment
Time (h)
Sort
ATAC-seq
mock
2
8
24
48
all
HIV-1-GFP
GFP-neg
GFP-pos
CD86-low
CD86-high
ENCODE
NF-κB ChIP-seq
IRF3 ChIP-seq
STAT1 ChIP-seq
CD14+ rep 1
CD14+ rep 2
D
IFNB1
IFNL1
CXCL10
CXCL11
ISG15
LY6E
USP18
IFIT1

**Figure S2. Time-dependent chromatin opening at IFN and ISG promoters occurs during HIV-1-GFP infection in DCs. Related to Figure 2.**

(A) PCA plots for ATAC-seq data from DCs that were either mock treated or infected with HIV-1-GFP. All time points were grouped by condition (shades of blue or red for mock and HIV-1-GFP, respectively) and contour lines extend outward from the condition mean to mark standard deviations. Marginal curves show the density of sample position for each donor.

(B) PCA plots for ATAC-seq samples as in (A) showing mean-centered contour lines for conditions separated by time (2, 8, 24, and 48 h). Color scale bars indicate mock, HIV-1-GFP, and specific populations.

(C-D) ATAC-seq tracks at the indicated gene start sites aligned with tracks from ENCODE ChIP-seq for NF-κB, STAT1, IRF3, and DNAse hypersensitivity data from independent CD14+ monocyte samples. Genes with accessible chromatin at baseline are shown in (C) compared to baseline inaccessible chromatin (D). ENCODE data is scaled equally for (C) and (D). ATAC-seq tracks represent merged files across 3 biological replicates and were visualized using IGV.

Figure S3

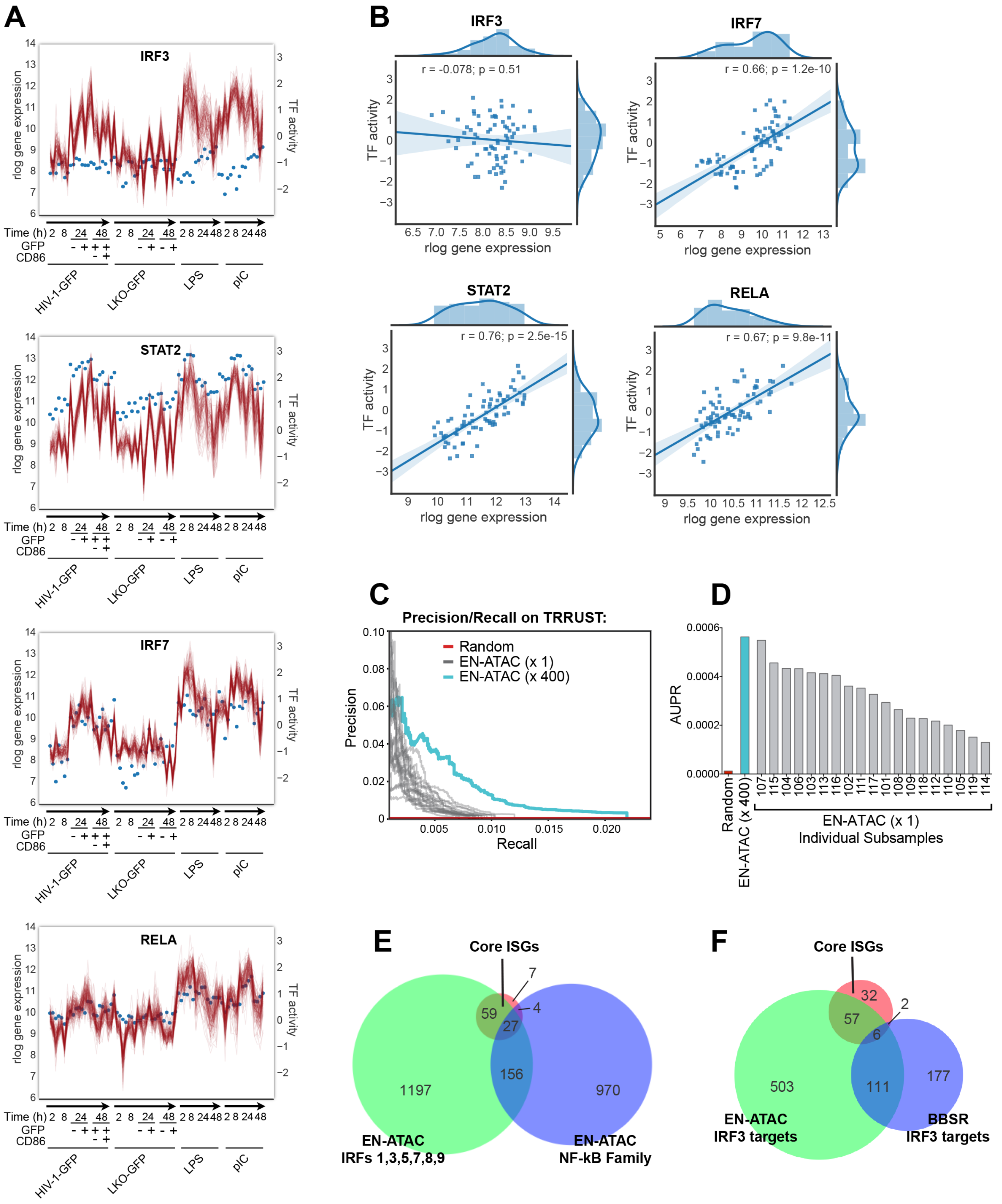
A
IRF3
STAT2
IRF7
RELA
rlog gene expression
TF activity
Time (h) 2 8 24 48 2 8 24 48 2 8 24 48 2 8 24 48
GFP - + + + - + - +
CD86 - +
HIV-1-GFP
LKO-GFP
LPS
pIC
B
IRF3
r = -0.078; p = 0.51
IRF7
r = 0.66; p = 1.2e-10
STAT2
r = 0.76; p = 2.5e-15
RELA
r = 0.67; p = 9.8e-11
TF activity
rlog gene expression
C
Precision/Recall on TRRUST:
Random
EN-ATAC (x 1)
EN-ATAC (x 400)
Precision
Recall
D
AUPR
Random
EN-ATAC (x 400)
107 115 104 106 103 113 116 102 111 117 101 108 109 118 112 110 105 119 114
EN-ATAC (x 1)
Individual Subsamples
E
Core ISGs
59
7
4
27
156
1197
970
EN-ATAC
IRFs 1,3,5,7,8,9
EN-ATAC
NF-kB Family
F
Core ISGs
32
57
2
6
503
111
177
EN-ATAC
IRF3 targets
BBSR
IRF3 targets

**Figure S3. TF activity estimates combined with multiple subsampling of prior information improves network inference. Related to Figure 3.**

(A) Joint plots for gene expression and estimated activities for the indicated TFs. Blue points are scaled to left y-axis and show rlog gene expression from 4 unique donors in series. 100 individual activity estimates are displayed as red lines and scaled on the right y-axis as normalized TF activity. The x-axis indicates time and stimulation condition.

(B) Pearson correlation plots showing the relationship between estimated activity and rlog gene expression for the TFs shown in (A).

(C) Precision/Recall performance measured against the TRRUST database for the final EN-ATAC (x 400) network (cyan) compared to 19 individual network runs generated from single subsamples of the prior (grey) and a network generated from random (red) (see STAR Methods).

(D) AUPR curves for the networks shown as in (C) scored against the TRRUST database.

(E) Proportional Venn diagrams displaying overlap between targets predicted in the final EN-ATAC network for the indicated IRF family members, NF-κB family members, and 97 “core” mammalian ISGs (Shaw et al., 2017).

(F) Proportional Venn diagrams displaying overlap between targets of IRF3 predicted by the EN-ATAC network, IRF3 targets predicted by BBSR, and core mammalian ISGs.

**Figure S4**

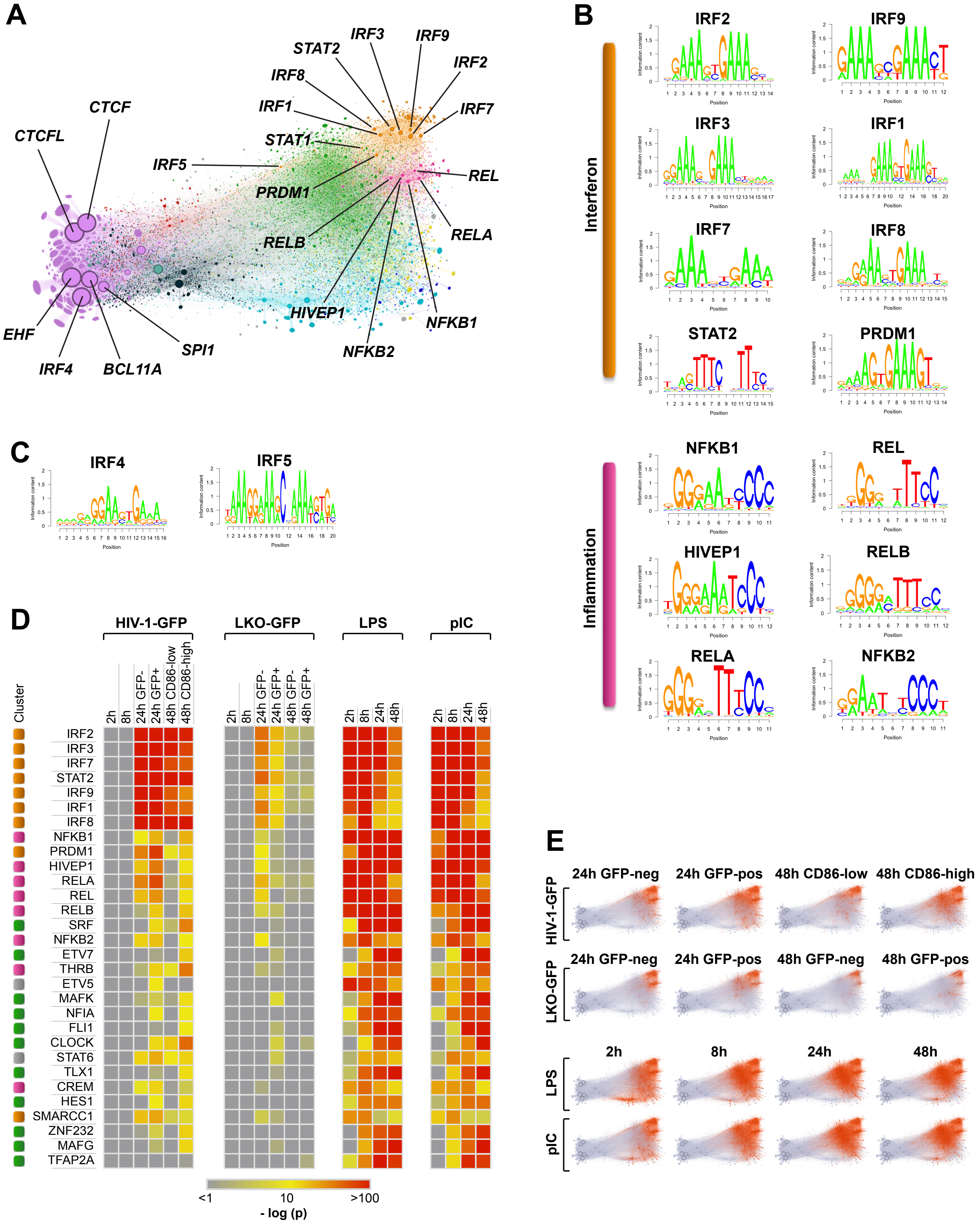
A
CTCFL
CTCF
IRF5
IRF8
STAT2
IRF3
IRF9
IRF2
IRF1
IRF7
STAT1
PRDM1
REL
RELB
RELA
HIVEP1
NFKB1
NFKB2
EHF
IRF4
BCL11A
SPI1
B
Interferon
IRF2
IRF9
IRF3
IRF1
IRF7
IRF8
STAT2
PRDM1
Inflammation
NFKB1
REL
HIVEP1
RELB
RELA
NFKB2
Information content
Position
C
IRF4
IRF5
D
Cluster
HIV-1-GFP
LKO-GFP
LPS
pIC
2h
8h
24h GFP-
24h GFP+
48h CD86-low
48h CD86-high
48h GFP-
48h GFP+
24h
48h
IRF2
IRF3
IRF7
STAT2
IRF9
IRF1
IRF8
NFKB1
PRDM1
HIVEP1
RELA
REL
RELB
SRF
NFKB2
ETV7
THRB
ETV5
MAFK
NFIA
FLI1
CLOCK
STAT6
TLX1
CREM
HES1
SMARCC1
ZNF232
MAFG
TFAP2A
<1
10
>100
- log (p)
E
HIV-1-GFP
LKO-GFP
LPS
pIC
24h GFP-neg
24h GFP-pos
48h CD86-low
48h CD86-high
48h GFP-neg
48h GFP-pos
2h
8h
24h
48h

**Figure S4. IRF and NF-κB family members populate dynamic areas of the network. Related to Figure 4.**

(A) Selected TF positions in the network as visualized by Gephi software. Edge line weights are reduced to accentuate nodes, with node size being relative to the number of direct connections. Color coding is consistent with cluster naming as in Figure 4.

(B) HOCOMOCO motif sequence logos for top TFs in Cluster 5 (IRF & Interferon) and Cluster 8 (NF-κB and Inflammation).

(C) HOCOMOCO motif sequence logos for IRF4 and IRF5

(D) Heat maps of hypergeometric tests for TF enrichment across all time series conditions for HIV-1-GFP, LKO-GFP, LPS, and polyI:C as compared to mock treatment. TFs were ranked by their weighted z-scores combined across all contrasts. Data is displayed as -log (p-value).

(E) Network activity across the time series shown for HIV-1-GFP, LKO-GFP, LPS, and polyI:C as visualized by Gephi software. Red areas denote differential gene expression and track temporally with TF hypergeometric enrichment as shown in (D).

Figure S5

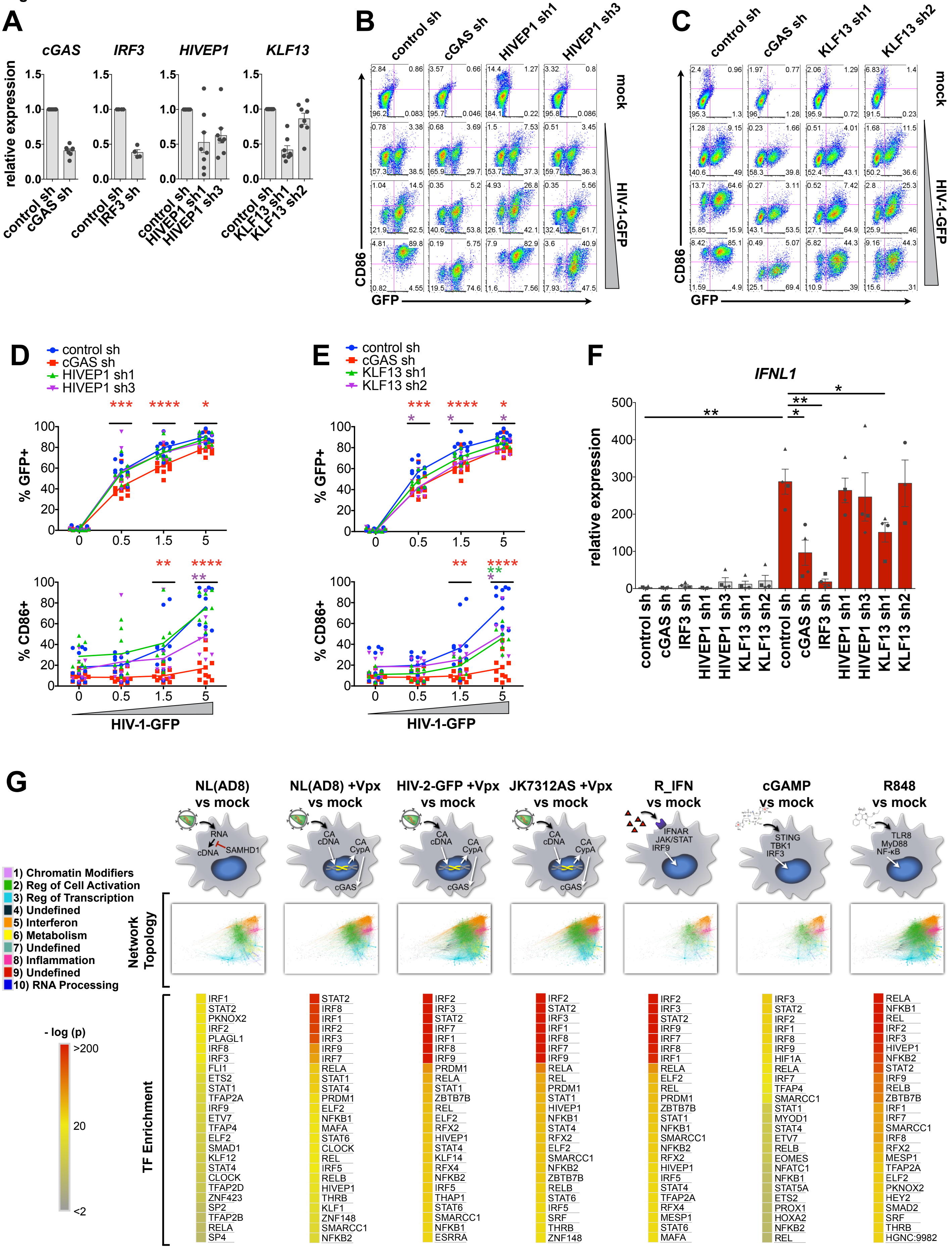

A
cGAS
IRF3
HIVEP1
KLF13
relative expression
control sh
cGAS sh
IRF3 sh
HIVEP1 sh1
HIVEP1 sh3
KLF13 sh1
KLF13 sh2
B
C
mock
HIV-1-GFP
CD86
GFP
D
E
% GFP+
% CD86+
HIV-1-GFP
F
IFNL1
G
NL(AD8) vs mock
NL(AD8) +Vpx vs mock
HIV-2-GFP +Vpx vs mock
JK7312AS +Vpx vs mock
R_IFN vs mock
cGAMP vs mock
R848 vs mock
RNA
cDNA
SAMHD1
CA
CypA
cGAS
IFNAR
JAK/STAT
IRF9
STING
TBK1
IRF3
TLR8
MyD88
NF-κB
1) Chromatin Modifiers
2) Reg of Cell Activation
3) Reg of Transcription
4) Undefined
5) Interferon
6) Metabolism
7) Undefined
8) Inflammation
9) Undefined
10) RNA Processing
Network Topology
TF Enrichment
- log (p)
>200
20
<2
IRF1 STAT2 PKNOX2 IRF2 PLAGL1 IRF8 IRF3 FLI1 ETS2 STAT1 TFAP2A IRF9 ETV7 TFAP4 ELF2 SMAD1 KLF12 STAT4 CLOCK TFAP2D ZNF423 SP2 TFAP2B RELA SP4
STAT2 IRF8 IRF1 IRF2 IRF3 IRF9 IRF7 RELA STAT1 STAT4 PRDM1 ELF2 NFKB1 MAFA STAT6 CLOCK REL IRF5 RELB HIVEP1 THRB KLF1 ZNF148 SMARCC1 NFKB2
IRF2 IRF3 STAT2 IRF7 IRF1 IRF8 IRF9 PRDM1 RELA STAT1 ZBTB7B REL ELF2 RFX2 HIVEP1 STAT4 KLF14 RFX4 NFKB2 IRF5 THAP1 STAT6 SMARCC1 NFKB1 ESRRA
IRF2 STAT2 IRF3 IRF1 IRF8 IRF7 IRF9 RELA REL PRDM1 STAT1 HIVEP1 NFKB1 STAT4 RFX2 ELF2 SMARCC1 NFKB2 ZBTB7B RELB STAT6 IRF5 SRF THRB ZNF148
IRF2 IRF3 STAT2 IRF9 IRF7 IRF8 IRF1 RELA ELF2 REL PRDM1 ZBTB7B STAT1 NFKB1 SMARCC1 NFKB2 RFX2 HIVEP1 IRF5 STAT4 TFAP2A RFX4 MESP1 STAT6 MAFA
IRF3 STAT2 IRF2 IRF1 IRF8 IRF9 HIF1A RELA IRF7 TFAP4 SMARCC1 STAT1 MYOD1 STAT4 ETV7 RELB EOMES NFATC1 NFKB1 STAT5A ETS2 PROX1 HOXA2 NFKB2 REL
RELA NFKB1 REL IRF2 IRF3 HIVEP1 NFKB2 STAT2 IRF9 RELB ZBTB7B IRF1 IRF7 SMARCC1 IRF8 RFX2 MESP1 TFAP2A ELF2 PKNOX2 HEY2 SMAD2 SRF THRB HGNC:9982

**Figure S5. Candidate screening of predicted IFN regulators reveal positive- and negative-acting factors in HIV-driven innate responses. Related to Figure 5.**

(A) qPCR assessment of target knockdown in lenti-shRNA modified DCs at day 5 after transduction. n = 4-8 donors

(B-C) Flow cytometry of CD86 vs GFP expression in shRNA-modified DCs 48 h after mock treatment or infection with HIV-1-GFP.

(D-E) Plots showing %GFP+ and %CD86+ from DCs treated as in (B-C). Plots represent pooled data from multiple donors. Identical data points for control and cGAS sh conditions are shown on panels D & E for comparison to HIVEP1 and KLF13 shRNA treatments, respectively. n = 11 (control sh); 11 (cGAS sh); HIVEP1 sh1/3 (8); KLF13 sh1/2 (7).

(F) qPCR of *IFNL1* gene expression in DCs for the indicated shRNA conditions that were mock treated or infected with HIV-1-GFP for 24 h (MOI 2). n = 4 for all conditions except KLF13 sh2.

(G) Illustration of additional viral perturbations and classic innate agonists that were incorporated into the network. Differential gene expression for each indicated contrast was visualized by Gephi software and color-coded by network cluster as shown (Network Topology). Heat maps of hypergeometric test results for each contrast are also shown (TF enrichment).

For (D-F), * $p < 0.05$; ** $p < 0.01$; *** $p < 0.001$. Data represent mean ± SEM.

Figure S6

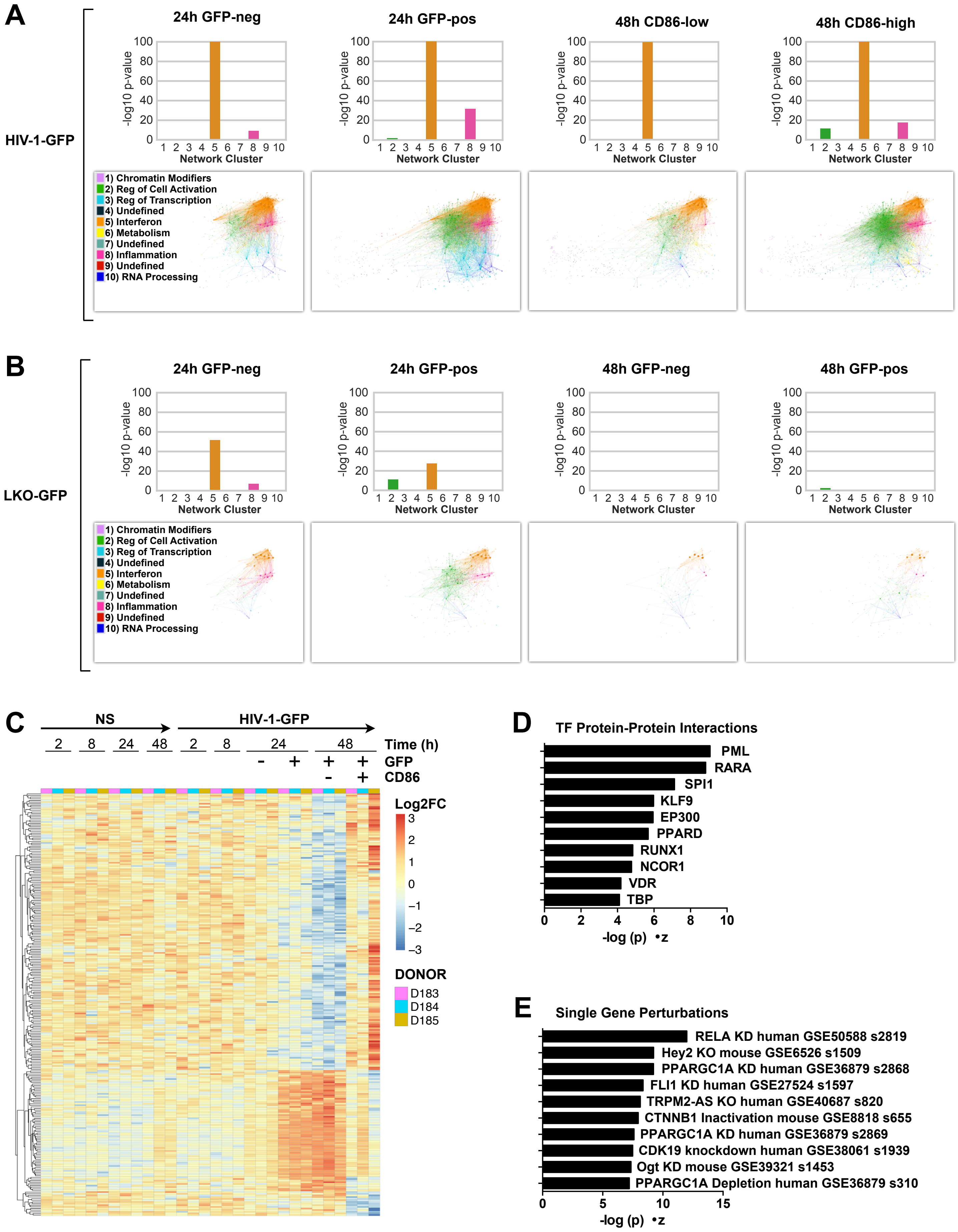

A
HIV-1-GFP
24h GFP-neg
24h GFP-pos
48h CD86-low
48h CD86-high
-log10 p-value
Network Cluster
1) Chromatin Modifiers
2) Reg of Cell Activation
3) Reg of Transcription
4) Undefined
5) Interferon
6) Metabolism
7) Undefined
8) Inflammation
9) Undefined
10) RNA Processing
B
LKO-GFP
24h GFP-neg
24h GFP-pos
48h GFP-neg
48h GFP-pos
C
NS
HIV-1-GFP
Time (h)
GFP
CD86
Log2FC
DONOR
D183
D184
D185
D
TF Protein-Protein Interactions
PML
RARA
SPI1
KLF9
EP300
PPARD
RUNX1
NCOR1
VDR
TBP
-log (p) •z
E
Single Gene Perturbations
RELA KD human GSE50588 s2819
Hey2 KO mouse GSE6526 s1509
PPARGC1A KD human GSE36879 s2868
FLI1 KD human GSE27524 s1597
TRPM2-AS KO human GSE40687 s820
CTNNB1 Inactivation mouse GSE8818 s655
PPARGC1A KD human GSE36879 s2869
CDK19 knockdown human GSE38061 s1939
Ogt KD mouse GSE39321 s1453
PPARGC1A Depletion human GSE36879 s310
-log (p) •z

**Figure S6. DE gene enrichment by cluster and changes in open chromatin between mature and immature DCs predict factors regulating cell activation. Related to Figure 6.**

(A) Enrichment of differentially expressed genes separated by network clusters for the following contrasts compared to mock: 24 h HIV-1-GFP GFP-positive; 24 h HIV-1-GFP GFP-negative; 48 h HIV-1-GFP CD86-low; 48 h HIV-1-GFP CD86-high. Differential gene expression for each indicated contrast is visualized by Gephi software and color-coded by network cluster as shown.

(B) Similar to (A) except for the indicated contrasts compared to mock for LKO-GFP.

(C) Heat map showing genes associated with differentially accessible chromatin identified from ATAC-seq analysis for 48 h HIV-1-GFP CD86-high compared to CD86-low sorted populations. Peaks were considered differential at FDR < 0.1.

(D-E) Potential Protein-Protein interactions (D) and Single Gene Perturbations (E) predicted from pathway analysis of genes linked with differentially accessible chromatin for the CD86-high vs CD86-low contrast.

Figure S7

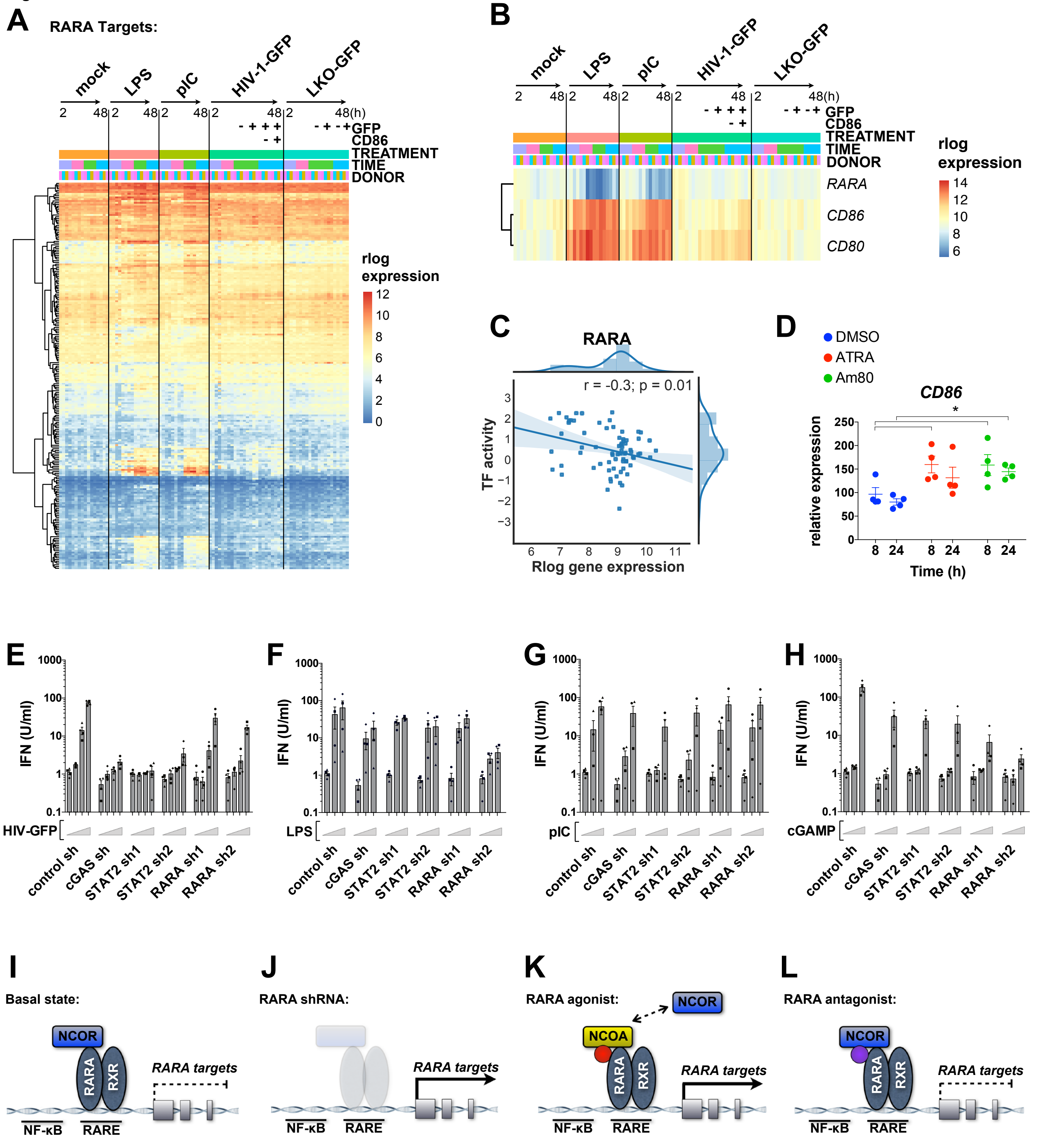
A
RARA Targets:
mock
LPS
pIC
HIV-1-GFP
LKO-GFP
2 48 2 48 2 48 2 48 2 48(h)
GFP
CD86
TREATMENT
TIME
DONOR
rlog expression
12 10 8 6 4 2 0
B
RARA
CD86
CD80
rlog expression
14 12 10 8 6
C
RARA
r = -0.3; p = 0.01
TF activity
Rlog gene expression
D
DMSO
ATRA
Am80
CD86
relative expression
Time (h)
E
F
G
H
IFN (U/ml)
HIV-GFP
LPS
pIC
cGAMP
control sh
cGAS sh
STAT2 sh1
STAT2 sh2
RARA sh1
RARA sh2
I
Basal state:
J
RARA shRNA:
K
RARA agonist:
L
RARA antagonist:
NCOR
NCOA
RARA
RXR
RARA targets
NF-κB
RARE

**Figure S7. RARA perturbations diametrically increase inflammatory signaling and mitigate type I IFN responses. Related to Figure 7.**

(A) Heat map showing gene expression of RARA targets predicted in the EN-ATAC network (500k cutoff; positive beta values and outliers removed) for the time course and stimulations shown in DCs (LPS, pIC, HIV-1-GFP, and LKO-GFP at 2, 8, 24, and 48 h).

(B) Heat map of *RARA, CD86*, and *CD80* expression over the time course and stimulations shown as in (A).

(C) Scatter plot showing negative correlation between estimated TF activity and rlog gene expression for RARA across all time course RNA-seq samples.

(D) qPCR of *CD86* expression in DCs after challenge with the RARA agonists all-trans retinoic acid (ATRA) and Am80 compared to vehicle alone (DMSO).

(E-H) Reporter bioassay of type I IFN activity in DC supernatants after stimulation. For (E), HIV-1-GFP MOI = 0, 1.5, 5. For (F) LPS = 0, 3, 10 ng/ml. For (G), polyI:C = 0, 1, 3 μg/ml. For (H), cGAMP = 0, 1, 5 μg/ml.

(I-L) Mechanistic models for how RARA perturbation influences gene expression.

For (D-H), * $p < 0.05$. Data represent mean ± SEM.

**Table S1. Metadata. Related to Figure 1.**

| SAMPLE | TIME | TREATMENT_CODE | DONOR_CODE | NETWORK_NORMALIZATION_GROU | VPX | OUTLIER | INCLUDE_IN_ANALYSIS |
|---|---|---|---|---|---|---|---|
| SL43835 | 0 | NS | D493 | CONTROL | NA | N | 1 |
| SL43833 | 0 | NS | D50 | CONTROL | NA | N | 1 |
| SL43834 | 0 | NS | D51 | CONTROL | NA | N | 1 |
| SL20541 | 2 | NS | 223 | CONTROL | NA | N | 1 |
| SL19557 | 2 | NS | 225 | CONTROL | NA | N | 1 |
| SL34329 | 2 | NS | 234 | CONTROL | NA | N | 1 |
| SL34334 | 2 | NS | 235 | CONTROL | NA | N | 1 |
| SL20542 | 2 | lps | 223 | SL20542 | NA | N | 1 |
| SL19558 | 2 | lps | 225 | SL19558 | NA | N | 1 |
| SL34330 | 2 | lps | 234 | SL34330 | NA | Y | 0 |
| SL34335 | 2 | lps | 235 | SL34335 | NA | Y | 0 |
| SL20543 | 2 | poly iC | 223 | SL20543 | NA | N | 1 |
| SL19559 | 2 | poly iC | 225 | SL19559 | NA | N | 1 |
| SL34331 | 2 | poly iC | 234 | SL34331 | NA | N | 1 |
| SL34336 | 2 | poly iC | 235 | SL34336 | NA | N | 1 |
| SL20544 | 2 | (G)HIVGFP + VPX | 223 | SL20544 | SIV_VPX | N | 1 |
| SL19560 | 2 | (G)HIVGFP + VPX | 225 | SL19560 | SIV_VPX | N | 1 |
| SL34332 | 2 | (G)HIVGFP + VPX | 234 | SL34332 | SIV_VPX | N | 1 |
| SL34337 | 2 | (G)HIVGFP + VPX | 235 | SL34337 | SIV_VPX | Y | 0 |
| SL19556 | 2 | (G)plko.1GFP + VPX | 223 | SL19556 | SIV_VPX | N | 1 |
| SL19561 | 2 | (G)plko.1GFP + VPX | 225 | SL19561 | SIV_VPX | N | 1 |
| SL34333 | 2 | (G)plko.1GFP + VPX | 234 | SL34333 | SIV_VPX | N | 1 |
| SL34338 | 2 | (G)plko.1GFP + VPX | 235 | SL34338 | SIV_VPX | N | 1 |
| SL45362 | 6 | cGAMP_lipo | D493 | SL45362 | NA | N | 1 |
| SL45361 | 6 | cGAMP_lipo | D50 | SL45361 | NA | N | 1 |
| SL46824 | 6 | cGAMP_lipo | D51 | SL46824 | NA | N | 1 |
| SL46823 | 6 | pdCdG_lipo | D493 | SL46823 | NA | N | 1 |
| SL46821 | 6 | pdCdG_lipo | D50 | SL46821 | NA | N | 1 |
| SL46822 | 6 | pdCdG_lipo | D51 | SL46822 | NA | N | 1 |
| SL46820 | 6 | pIC | D493 | SL46820 | NA | N | 1 |
| SL46818 | 6 | pIC | D50 | SL46818 | NA | N | 1 |
| SL46819 | 6 | pIC | D51 | SL46819 | NA | N | 1 |
| SL45364 | 6 | R848 | D493 | SL45364 | NA | N | 1 |
| SL46825 | 6 | R848 | D50 | SL46825 | NA | N | 1 |
| SL45363 | 6 | R848 | D51 | SL45363 | NA | N | 1 |
| SL45366 | 6 | recombinant_IFN | D493 | SL45366 | NA | N | 1 |
| SL45365 | 6 | recombinant_IFN | D50 | SL45365 | NA | N | 1 |
| SL46826 | 6 | recombinant_IFN | D51 | SL46826 | NA | N | 1 |
| SL19562 | 8 | NS | 223 | CONTROL | NA | N | 1 |
| SL19567 | 8 | NS | 225 | CONTROL | NA | N | 1 |
| SL34339 | 8 | NS | 234 | CONTROL | NA | N | 1 |
| SL34574 | 8 | NS | 235 | CONTROL | NA | N | 1 |
| SL19563 | 8 | lps | 223 | SL19563 | NA | N | 1 |
| SL19568 | 8 | lps | 225 | SL19568 | NA | N | 1 |
| SL34340 | 8 | lps | 234 | SL34340 | NA | N | 1 |
| SL34575 | 8 | lps | 235 | SL34575 | NA | N | 1 |
| SL19564 | 8 | poly iC | 223 | SL19564 | NA | N | 1 |
| SL19569 | 8 | poly iC | 225 | SL19569 | NA | N | 1 |
| SL34571 | 8 | poly iC | 234 | SL34571 | NA | N | 1 |
| SL34576 | 8 | poly iC | 235 | SL34576 | NA | N | 1 |
| SL19565 | 8 | (G)HIVGFP + VPX | 223 | SL19565 | SIV_VPX | N | 1 |
| SL19570 | 8 | (G)HIVGFP + VPX | 225 | SL19570 | SIV_VPX | N | 1 |
| SL34572 | 8 | (G)HIVGFP + VPX | 234 | SL34572 | SIV_VPX | N | 1 |
| SL34577 | 8 | (G)HIVGFP + VPX | 235 | SL34577 | SIV_VPX | N | 1 |
| SL19566 | 8 | (G)plko.1GFP + VPX | 223 | SL19566 | SIV_VPX | N | 1 |
| SL19571 | 8 | (G)plko.1GFP + VPX | 225 | SL19571 | SIV_VPX | N | 1 |
| SL34573 | 8 | (G)plko.1GFP + VPX | 234 | SL34573 | SIV_VPX | N | 1 |
| SL34578 | 8 | (G)plko.1GFP + VPX | 235 | SL34578 | SIV_VPX | N | 1 |
| SL19572 | 24 | NS | 223 | CONTROL | NA | N | 1 |
| SL19579 | 24 | NS | 225 | CONTROL | NA | N | 1 |
| SL19600 | 24 | NS | 234 | CONTROL | NA | N | 1 |
| SL19607 | 24 | NS | 235 | CONTROL | NA | N | 1 |
| SL19573 | 24 | lps | 223 | SL19573 | NA | N | 1 |
| SL19580 | 24 | lps | 225 | SL19580 | NA | N | 1 |
| SL19601 | 24 | lps | 234 | SL19601 | NA | N | 1 |
| SL19608 | 24 | lps | 235 | SL19608 | NA | N | 1 |
| SL19574 | 24 | poly iC | 223 | SL19574 | NA | N | 1 |
| SL19581 | 24 | poly iC | 225 | SL19581 | NA | N | 1 |
| SL19602 | 24 | poly iC | 234 | SL19602 | NA | N | 1 |
| SL19609 | 24 | poly iC | 235 | SL19609 | NA | N | 1 |
| SL19575 | 24 | GFP- (G)HIVGFP + VPX | 223 | SL19575 | SIV_VPX | N | 1 |
| SL19582 | 24 | GFP- (G)HIVGFP + VPX | 225 | SL19582 | SIV_VPX | N | 1 |
| SL19604 | 24 | GFP- (G)HIVGFP + VPX | 234 | SL19604 | SIV_VPX | N | 1 |
| SL19611 | 24 | GFP- (G)HIVGFP + VPX | 235 | SL19611 | SIV_VPX | N | 1 |
| SL19576 | 24 | GFP+ (G)HIVGFP + VPX | 223 | SL19576 | SIV_VPX | N | 1 |
| SL19583 | 24 | GFP+ (G)HIVGFP + VPX | 225 | SL19583 | SIV_VPX | N | 1 |
| SL19603 | 24 | GFP+ (G)HIVGFP + VPX | 234 | SL19603 | SIV_VPX | N | 1 |
| SL19610 | 24 | GFP+ (G)HIVGFP + VPX | 235 | SL19610 | SIV_VPX | N | 1 |
| SL19577 | 24 | GFP- (G)plko.1GFP + VPX | 223 | SL19577 | SIV_VPX | N | 1 |
| SL19584 | 24 | GFP- (G)plko.1GFP + VPX | 225 | SL19584 | SIV_VPX | N | 1 |
| SL19606 | 24 | GFP- (G)plko.1GFP + VPX | 234 | SL19606 | SIV_VPX | N | 1 |
| SL19613 | 24 | GFP- (G)plko.1GFP + VPX | 235 | SL19613 | SIV_VPX | N | 1 |
| SL19578 | 24 | GFP+ (G)plko.1GFP + VPX | 223 | SL19578 | SIV_VPX | N | 1 |

| | | | | | | | |
|---|---|---|---|---|---|---|---|
| SL19585 | 24 | GFP+ (G)plko.1GFP + VPX | 225 | SL19585 | SIV_VPX | N | 1 |
| SL19605 | 24 | GFP+ (G)plko.1GFP + VPX | 234 | SL19605 | SIV_VPX | N | 1 |
| SL19612 | 24 | GFP+ (G)plko.1GFP + VPX | 235 | SL19612 | SIV_VPX | N | 1 |
| SL43844 | 24 | HIV2_P86HA | D493 | SL43844 | NA | N | 1 |
| SL43842 | 24 | HIV2_P86HA | D50 | SL43842 | NA | N | 1 |
| SL43843 | 24 | HIV2_P86HA | D51 | SL43843 | NA | N | 1 |
| SL45353 | 24 | HIV2_P86HA+CsA | D493 | SL45353 | NA | N | 1 |
| SL45351 | 24 | HIV2_P86HA+CsA | D50 | SL45351 | NA | N | 1 |
| SL45352 | 24 | HIV2_P86HA+CsA | D51 | SL45352 | NA | N | 1 |
| SL45355 | 24 | HIV2_WT | D493 | SL45355 | NA | N | 1 |
| SL45354 | 24 | HIV2_WT | D50 | SL45354 | NA | N | 1 |
| SL46813 | 24 | HIV2_WT | D51 | SL46813 | NA | N | 1 |
| SL46814 | 24 | JK7312AS | D493 | SL46814 | NA | N | 1 |
| SL45356 | 24 | JK7312AS | D50 | SL45356 | NA | N | 1 |
| SL45357 | 24 | JK7312AS | D51 | SL45357 | NA | N | 1 |
| SL46816 | 24 | NLAD8 | D493 | SL46816 | NA | N | 1 |
| SL45358 | 24 | NLAD8 | D50 | SL45358 | NA | N | 1 |
| SL46815 | 24 | NLAD8 | D51 | SL46815 | NA | N | 1 |
| SL45360 | 24 | NLAD8+SIV | D493 | SL45360 | SIV_VPX | N | 1 |
| SL46817 | 24 | NLAD8+SIV | D50 | SL46817 | SIV_VPX | N | 1 |
| SL45359 | 24 | NLAD8+SIV | D51 | SL45359 | SIV_VPX | N | 1 |
| SL43838 | 24 | NS | D493 | CONTROL | NA | N | 1 |
| SL43836 | 24 | NS | D50 | CONTROL | NA | N | 1 |
| SL43837 | 24 | NS | D51 | CONTROL | NA | N | 1 |
| SL43841 | 24 | NS+CsA | D493 | SL43841 | NA | N | 1 |
| SL43839 | 24 | NS+CsA | D50 | SL43839 | NA | N | 1 |
| SL43840 | 24 | NS+CsA | D51 | SL43840 | NA | N | 1 |
| SL13423 | 48 | NS | 180 | CONTROL | NA | N | 1 |
| SL11695 | 48 | (G)HIVGFP+ Vpx | 179 | SL11695 | SIV_VPX | Y | 1 |
| SL12686 | 48 | (G)HIVGFP+ Vpx | 179 | SL12686 | SIV_VPX | Y | 1 |
| SL12692 | 48 | (G)HIVGFP+ Vpx | 180 | SL12692 | SIV_VPX | N | 1 |
| SL13422 | 48 | (G)HIVGFP+ Vpx | 180 | SL13422 | SIV_VPX | N | 1 |
| SL11697 | 48 | (G)HIVGFP+ Vpx(-) | 179 | SL11697 | SIV_VPX_NEG | N | 1 |
| SL12687 | 48 | (G)HIVGFP+ Vpx(-) | 179 | SL12687 | SIV_VPX_NEG | N | 1 |
| SL12693 | 48 | (G)HIVGFP+ Vpx(-) | 180 | SL12693 | SIV_VPX_NEG | N | 1 |
| SL13424 | 48 | (G)HIVGFP+ Vpx(-) | 180 | SL13424 | SIV_VPX_NEG | N | 1 |
| SL11696 | 48 | (G)HIVGFP | 179 | SL11696 | NA | N | 1 |
| SL11698 | 48 | (G)plk0.1GFP+Vpx | 179 | SL11698 | SIV_VPX | N | 1 |
| SL12688 | 48 | (G)plk0.1GFP+Vpx | 179 | SL12688 | SIV_VPX | N | 1 |
| SL12694 | 48 | (G)plk0.1GFP+Vpx | 180 | SL12694 | SIV_VPX | N | 1 |
| SL13425 | 48 | (G)plk0.1GFP+Vpx | 180 | SL13425 | SIV_VPX | Y | 1 |
| SL11700 | 48 | (G)plk0.1GFP+Vpx(-) | 179 | SL11700 | SIV_VPX_NEG | N | 1 |
| SL12689 | 48 | (G)plk0.1GFP+Vpx(-) | 179 | SL12689 | SIV_VPX_NEG | N | 1 |
| SL12695 | 48 | (G)plk0.1GFP+Vpx(-) | 180 | SL12695 | SIV_VPX_NEG | N | 1 |
| SL13427 | 48 | (G)plk0.1GFP+Vpx(-) | 180 | SL13427 | SIV_VPX_NEG | N | 1 |
| SL11699 | 48 | (G)plk0.1GFP | 179 | SL11699 | NA | N | 1 |
| SL13426 | 48 | (G)plk0.1GFP | 180 | SL13426 | NA | N | 1 |
| SL11701 | 48 | Vpx | 179 | SL11701 | SIV_VPX | N | 1 |
| SL12690 | 48 | Vpx | 179 | SL12690 | SIV_VPX | N | 1 |
| SL12696 | 48 | Vpx | 180 | SL12696 | SIV_VPX | N | 1 |
| SL13428 | 48 | Vpx | 180 | SL13428 | SIV_VPX | N | 1 |
| SL11702 | 48 | Vpx(-) | 179 | SL11702 | SIV_VPX_NEG | N | 1 |
| SL12691 | 48 | Vpx(-) | 179 | SL12691 | SIV_VPX_NEG | N | 1 |
| SL12697 | 48 | Vpx(-) | 180 | SL12697 | SIV_VPX_NEG | N | 1 |
| SL13429 | 48 | Vpx(-) | 180 | SL13429 | SIV_VPX_NEG | N | 1 |
| SL19586 | 48 | NS | 223 | CONTROL | NA | N | 1 |
| SL19593 | 48 | NS | 225 | CONTROL | NA | N | 1 |
| SL19614 | 48 | NS | 234 | CONTROL | NA | N | 1 |
| SL19621 | 48 | NS | 235 | CONTROL | NA | N | 1 |
| SL19587 | 48 | lps | 223 | SL19587 | NA | N | 1 |
| SL19594 | 48 | lps | 225 | SL19594 | NA | N | 1 |
| SL19615 | 48 | lps | 234 | SL19615 | NA | N | 1 |
| SL22753 | 48 | lps | 235 | SL22753 | NA | N | 1 |
| SL19588 | 48 | poly iC | 223 | SL19588 | NA | N | 1 |
| SL19595 | 48 | poly iC | 225 | SL19595 | NA | N | 1 |
| SL19616 | 48 | poly iC | 234 | SL19616 | NA | N | 1 |
| SL19622 | 48 | poly iC | 235 | SL19622 | NA | N | 1 |
| SL19589 | 48 | GFP+ 86- (G)HIVGFP + VF | 223 | SL19589 | SIV_VPX | N | 1 |
| SL19596 | 48 | GFP+ 86- (G)HIVGFP + VF | 225 | SL19596 | SIV_VPX | N | 1 |
| SL19618 | 48 | GFP+ 86- (G)HIVGFP + VF | 234 | SL19618 | SIV_VPX | N | 1 |
| SL19624 | 48 | GFP+ 86- (G)HIVGFP + VF | 235 | SL19624 | SIV_VPX | N | 1 |
| SL19590 | 48 | GFP+ 86+ (G)HIVGFP + VI | 223 | SL19590 | SIV_VPX | N | 1 |
| SL19597 | 48 | GFP+ 86+ (G)HIVGFP + VI | 225 | SL19597 | SIV_VPX | N | 1 |
| SL19617 | 48 | GFP+ 86+ (G)HIVGFP + VI | 234 | SL19617 | SIV_VPX | N | 1 |
| SL19623 | 48 | GFP+ 86+ (G)HIVGFP + VI | 235 | SL19623 | SIV_VPX | N | 1 |
| SL19599 | 48 | GFP- (G)plko.1GFP + VPX | 225 | SL19599 | SIV_VPX | N | 1 |
| SL19620 | 48 | GFP- (G)plko.1GFP + VPX | 234 | SL19620 | SIV_VPX | N | 1 |
| SL19598 | 48 | GFP+ (G)plko.1GFP + VPX | 225 | SL19598 | SIV_VPX | N | 1 |
| SL19619 | 48 | GFP+ (G)plko.1GFP + VPX | 234 | SL19619 | SIV_VPX | N | 1 |
| SL19625 | 48 | GFP+ (G)plko.1GFP + VPX | 235 | SL19625 | SIV_VPX | N | 1 |

### Table S7. shRNA and primer sequences. Related to STAR Methods.

| shRNA clone name | TRCN | sequence | Source or Sigma validation in % knockdown |
|---|---|---|---|
| control sh | **N/A** | CCGGCAACAAGATGAAGAGCACCAACTCGAGTTGGTGCTCTTCATCTTGTTGTTTTTG | |
| LacZ sh | **TRCN0000072229** | CCGGGCGATCGTAATCACCCGAGTGCTCGAGCACTCGGGTGATTACGATCGCTTTTTG | |
| NONO sh | **TRCN0000074562** | CCGGCAGGCGAAGTCTTCATTCATACTCGAGTATGAATGAAGACTTCGCCTGTTTTTG | 0.98 |
| cGAS sh | **TRCN0000149984** | CCGGCAACTACGACTAAAGCCATTTCTCGAGAAATGGCTTTAGTCGTAGTTGTTTTTTG | 0.85 |
| IRF3 sh | **N/A** | CCGGCTGCCTGGATGGCCAGTCACACCTGTGAAGCCACAGATGGGGTGTGACTGGCCATCCAGGCAGTTTTT | Johnson et al., 2018 |
| HIVEP1 sh1 | **TRCN0000235474** | CCGGTAGGCGTCTCAGTAGGTTTAACTCGAGTTAAACCTACTGAGACGCCTATTTTTG | 0.79 |
| HIVEP1 sh3 | **TRCN0000235476** | CCGGCATGCCGACCACAGGTTATTCCTCGAGGAATAACCTGTGGTCGGCATGTTTTTG | |
| KLF13 sh1 | **TRCN0000016923** | CCGGGCGAGAAAGTTTACGGGAAATCTCGAGATTTCCCGTAAACTTTCTCGCTTTTT | 0.93 |
| KLF13 sh2 | **TRCN0000016925** | CCGGCGGGCGAGAAGAAGTTCAGCTCTCGAGAGCTGAACTTCTTCTCGCCCGTTTTT | 0.85 |
| STAT2 sh1 | **TRCN0000364399** | CCGGTAGGACTGAGGATCCATTATTCTCGAGAATAATGGATCCTCAGTCCTATTTTTG | 0.88 |
| STAT2 sh2 | **TRCN0000364400** | CCGGTGTCTTCTGCTTCCGATATAACTCGAGTTATATCGGAAGCAGAAGACATTTTTG | 0.89 |
| RARA sh1 | **TRCN0000275554** | CCGGCAGCTTCCAGTTAGTGGATATCTCGAGATATCCACTAACTGGAAGCTGTTTTTG | 0.77 |
| RARA sh2 | **TRCN0000020370** | CCGGCCCAAGATGCTAATGAAGATTCTCGAGAATCTTCATTAGCATCTTGGGTTTTT | 0.8 |
| PRDM1 sh2 | **TRCN0000013609** | CCGGCGGGATGAACATCTACTTCTACTCGAGTAGAAGTAGATGTTCATCCCGTTTTT | |
| PRDM1 sh3 | **TRCN0000013610** | CCGGCGAAGCCATGAATCTCATTAACTCGAGTTAATGAGATTCATGGCTTCGTTTTT | |
| PRDM1 sh5 | **TRCN0000013612** | CCGGGCCTGAAAGTGTCTTTGCAAACTCGAGTTTGCAAAGACACTTTCAGGCTTTTT | |

| ATAC-seq primers | | | |
|---|---|---|---|
| Ad1_noMX | | AATGATACGGCGACCACCGAGATCTACACTCGTCGGCAGCGTCAGATGTG | Buenrostro et al., 2013 |
| Ad2.1_TAAGGCGA | | CAAGCAGAAGACGGCATACGAGATTCGCCTTAGTCTCGTGGGCTCGGAGATGT | Buenrostro et al., 2013 |
| Ad2.2_CGTACTAG | | CAAGCAGAAGACGGCATACGAGATCTAGTACGGTCTCGTGGGCTCGGAGATGT | Buenrostro et al., 2013 |
| Ad2.3_AGGCAGAA | | CAAGCAGAAGACGGCATACGAGATTTCTGCCTGTCTCGTGGGCTCGGAGATGT | Buenrostro et al., 2013 |
| Ad2.4_TCCTGAGC | | CAAGCAGAAGACGGCATACGAGATGCTCAGGAGTCTCGTGGGCTCGGAGATGT | Buenrostro et al., 2013 |
| Ad2.5_GGACTCCT | | CAAGCAGAAGACGGCATACGAGATAGGAGTCCGTCTCGTGGGCTCGGAGATGT | Buenrostro et al., 2013 |
| Ad2.6_TAGGCATG | | CAAGCAGAAGACGGCATACGAGATCATGCCTAGTCTCGTGGGCTCGGAGATGT | Buenrostro et al., 2013 |
| Ad2.7_CTCTCTAC | | CAAGCAGAAGACGGCATACGAGATGTAGAGAGGTCTCGTGGGCTCGGAGATGT | Buenrostro et al., 2013 |
| Ad2.8_CAGAGAGG | | CAAGCAGAAGACGGCATACGAGATCCTCTCTGGTCTCGTGGGCTCGGAGATGT | Buenrostro et al., 2013 |
| Ad2.9_GCTACGCT | | CAAGCAGAAGACGGCATACGAGATAGCGTAGCGTCTCGTGGGCTCGGAGATGT | Buenrostro et al., 2013 |
| Ad2.10_CGAGGCTG | | CAAGCAGAAGACGGCATACGAGATCAGCCTCGGTCTCGTGGGCTCGGAGATGT | Buenrostro et al., 2013 |
| Ad2.11_AAGAGGCA | | CAAGCAGAAGACGGCATACGAGATTGCCTCTTGTCTCGTGGGCTCGGAGATGT | Buenrostro et al., 2013 |
| Ad2.12_GTAGAGGA | | CAAGCAGAAGACGGCATACGAGATTCCTCTACGTCTCGTGGGCTCGGAGATGT | Buenrostro et al., 2013 |
| Ad2.13_GTCGTGAT | | CAAGCAGAAGACGGCATACGAGATATCACGACGTCTCGTGGGCTCGGAGATGT | Buenrostro et al., 2013 |
| Ad2.14_ACCACTGT | | CAAGCAGAAGACGGCATACGAGATACAGTGGTGTCTCGTGGGCTCGGAGATGT | Buenrostro et al., 2013 |
| Ad2.15_TGGATCTG | | CAAGCAGAAGACGGCATACGAGATCAGATCCAGTCTCGTGGGCTCGGAGATGT | Buenrostro et al., 2013 |
| Ad2.16_CCGTTTGT | | CAAGCAGAAGACGGCATACGAGATACAAACGGGTCTCGTGGGCTCGGAGATGT | Buenrostro et al., 2013 |
| Ad2.17_TGCTGGGT | | CAAGCAGAAGACGGCATACGAGATACCCAGCAGTCTCGTGGGCTCGGAGATGT | Buenrostro et al., 2013 |
| Ad2.18_GAGGGGTT | | CAAGCAGAAGACGGCATACGAGATAACCCCTCGTCTCGTGGGCTCGGAGATGT | Buenrostro et al., 2013 |
| Ad2.19_AGGTTGGG | | CAAGCAGAAGACGGCATACGAGATCCCAACCTGTCTCGTGGGCTCGGAGATGT | Buenrostro et al., 2013 |
| Ad2.20_GTGTGGTG | | CAAGCAGAAGACGGCATACGAGATCACCACACGTCTCGTGGGCTCGGAGATGT | Buenrostro et al., 2013 |
| Ad2.21_TGGGTTTC | | CAAGCAGAAGACGGCATACGAGATGAAACCCAGTCTCGTGGGCTCGGAGATGT | Buenrostro et al., 2013 |
| Ad2.22_TGGTCACA | | CAAGCAGAAGACGGCATACGAGATTGTGACCAGTCTCGTGGGCTCGGAGATGT | Buenrostro et al., 2013 |
| Ad2.23_TTGACCCT | | CAAGCAGAAGACGGCATACGAGATAGGGTCAAGTCTCGTGGGCTCGGAGATGT | Buenrostro et al., 2013 |
| Ad2.24_CCACTCCT | | CAAGCAGAAGACGGCATACGAGATAGGAGTGGGTCTCGTGGGCTCGGAGATGT | Buenrostro et al., 2013 |
